# CsrA Shows Selective Regulation of sRNA-mRNA Networks

**DOI:** 10.1101/2023.03.29.534774

**Authors:** Alejandra Matsuri Rojano-Nisimura, Trevor R. Simmons, Abigail N. Leistra, Mia K. Mihailovic, Ryan Buchser, Alyssa M. Ekdahl, Isabella Joseph, Nicholas C. Curtis, Lydia M. Contreras

## Abstract

Post-transcriptional regulation, by small RNAs (sRNAs) as well as the global Carbon Storage Regulator A (CsrA) protein, play critical roles in bacterial metabolic control and stress responses. The CsrA protein affects selective sRNA-mRNA networks, in addition to regulating transcription factors and sigma factors, providing additional avenues of cross talk between other stress-response regulators. Here, we expand the known set of sRNA-CsrA interactions and study their regulatory effects. *In vitro* binding assays confirm novel CsrA interactions with ten sRNAs, many of which are previously recognized as key regulatory nodes. Of those 10 sRNA, we identify that McaS, FnrS, SgrS, MicL, and Spot42 interact with CsrA *in vivo*. We find that the presence of CsrA impacts the downstream regulation of mRNA targets of the respective sRNA. *In vivo* evidence supports enhanced CsrA McaS-*csgD* mRNA repression and showcase CsrA-dependent repression of the *fucP* mRNA via the Spot42 sRNA. We additionally identify SgrS and FnrS as potential new sRNA sponges of CsrA. Overall, our results further support the expanding impact of the Csr system on cellular physiology via CsrA impact on the regulatory roles of these sRNAs.

## Introduction

Non-coding RNAs have emerged as potent regulators of gene expression in cellular metabolism and stress responses (**Gottesman et al., 2006; Leistra et al., 2019; Vazquez-Anderson & Contreras, 2013; Wagner & Romby, 2015**). In bacteria, one type of regulatory non-coding RNAs, small RNAs (sRNAs), bind and affect the expression of their target genes at the protein and RNA level. For example, antisense sRNAs regulate a target mRNA via an extended region of perfectly complementary nucleotides to alter translation or stability of the mRNA (**Hör & Matera, 2020**). Alternatively, trans-acting sRNAs, oftentimes encoded in intergenic regions in the DNA, recognize one or more target mRNAs via regions of 15-40 nucleotides of limited complementarity (**Gottesman & Storz, 2011; Villa et al., 2017**). These interactions result in a diversity of regulatory outcomes and mechanisms (reviewed in **De Lay et al., 2013; Hör J Matera G, 2020; Nitzan et al., 2017; Wagner & Romby, 2015**).

Due to the vast regulatory functions of sRNAs, many efforts have been made to elucidate native regulatory roles of individual sRNAs **(Beisel & Storz, 2011; Durand & Storz, 2010; Guo et al., 2014; Lalaouna et al., 2015; Thomason et al., 2012; Vanderpool & Gottesman, 2004).** In *E. coli,* almost 100 sRNAs have been confirmed, targeting nearly 70 unique mRNAs, with 13 of these mRNAs regulated by multiple sRNAs **(Hör J Matera G, 2020; Mihailovic et al., 2018; J. Wang et al., 2016**). In this paper, we define a confirmed mRNA target as one in which a regulatory outcome and binding interface has been determined experimentally (*i.e.*, reporter assays with compensatory mutations or *in vitro* binding assays). However, outside of this confirmed mRNA target pool, a much broader set of putative sRNA-mRNA interactions have been inferred from recent high-throughput studies (**Iosub et al., 2020; Melamed et al., 2016, 2020; Waters et al., 2017**). A complex, dynamic network of stress response and metabolic control emerges from these data (Figure 1A) (**Nitzan et al., 2017; Wagner & Romby, 2015**), that is believed, in many cases, to hinge on the Hfq chaperone protein (**Kavita et al., 2018; Vogel & Luisi, 2011**).

**Figure 1.**
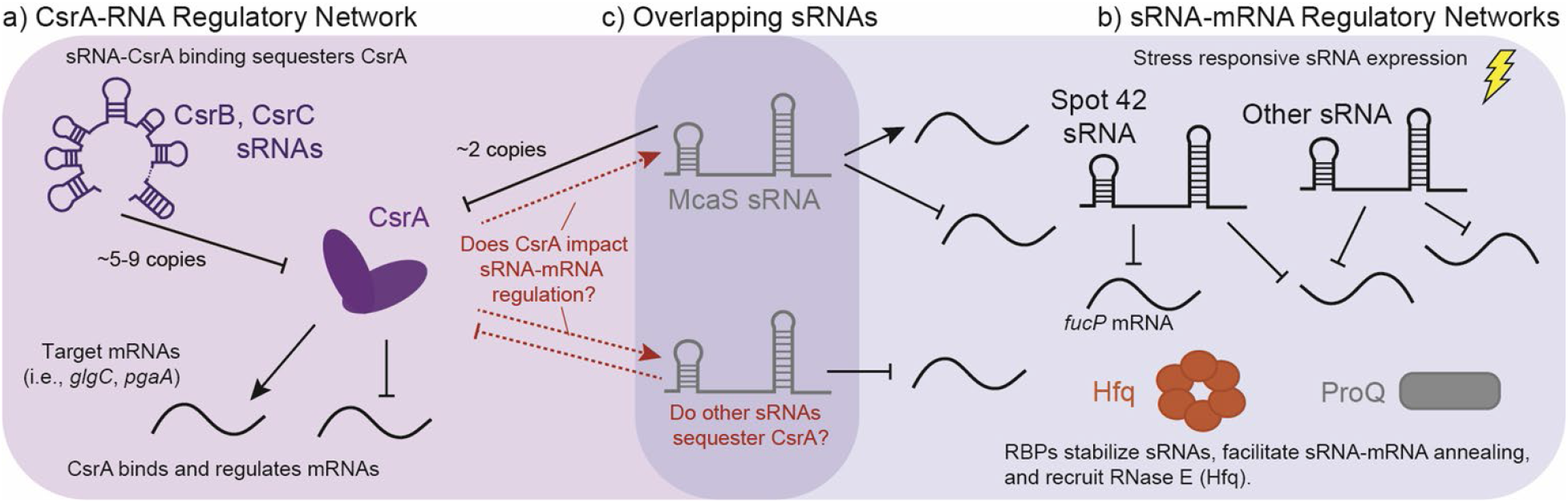
Overlap of sRNA-mRNA and CsrA-RNA post-transcriptional regulatory networks. 1a) The canonical CsrA regulatory network: CsrA binds mRNA targets to either repress or active translation. CsrA is then regulated by sRNAs CsrB and CsrB, which can sequester up to 9 and 5 copies of CsrA, respectively. 1b) Concurrently, sRNAs regulate mRNAs targets in response to external stimuli, these interactions are assisted by RNA Binding Proteins (RBPs) such as Hfq and ProQ. 1c) sRNAs such as McaS are overlapping sRNAs that both regulate mRNA targets, as well as CsrA. There is potentially many other sRNAs that can fall into both regulatory modes.

The hexameric Hfq protein is a global RNA-Binding Protein (RBP) that binds trans-acting sRNAs and their target mRNAs at well-characterized sequence motifs to facilitate inter-molecular base pairing **(Holmqvist & Vogel, 2018)**. Several explanations have been proposed for the RNA chaperone function of Hfq, including but not limited to: (i) stabilization of the sRNA, (ii) facilitation of inter-molecular base pairing by unfolding RNA secondary structures, and (iii) increase of the local sRNA concentration to augment mRNA pairing and regulation probability (**Santiago-Frangos and Woodson, 2018; Vogel and Luisi, 2011; Wagner et al., 2013**). However, the specific role of Hfq in each sRNA-mRNA interaction seems to remain variable, with many sRNAs sufficient to disrupt mRNA translation without Hfq, predominantly when the sRNA directly occludes the ribosome binding site of the respective mRNA (**Maki et al., 2008; Morita et al. 2006; Prevost et al., 2011**). For Hfq-dependent sRNAs, it has been shown that some sRNAs compete with each other for binding sites under Hfq-limited cellular conditions (**Santiago-Frangos et al., 2016**), suggesting complex and dynamic global post-transcriptional networks.

Despite the extensive regulatory role of Hfq across bacterial post-transcriptional networks, this protein is not the only global RBP that influences sRNA-mediated post-transcriptional regulation. More recently, the FinO-like protein ProQ was discovered as a global RBP through Grad-Seq (Figure. 1A). Like Hfq, ProQ has been shown to facilitate sRNA-mRNA interactions directly (**Holmqvist et al., 2018; Smirnov et al. 2016, 2017**), although using structural-specificity as opposed to sequence-specificity. RIL-Seq results show that in *E. coli,* ProQ and Hfq have approximately 100 shared target sRNA-mRNA pairs (**Melamed et al., 2020**). However, some organisms with documented trans-acting sRNAs-mRNA target pairs lack both Hfq and ProQ altogether; some examples include gram positive bacteria *Deinococcus radiodurans* (**Tsai et al., 2015; Villa et al. 2021)** and *Mycobacterium tuberculosis* (**Dichiara et al. 2010, Gerrick et al., 2018; Taneja & Dutta, 2019)**. In such organisms, KH domain proteins have started to gain attention as potential matchmaking RBPs (**Olejniczak et al., 2022**) due to their ability to associate with sRNAs (**Hör et al., 2020; Lamm-Schmidt et al., 2021)**. This reinforces the notion that other RBPs may be involved in sRNA-mRNA regulation (**Haning et al., 2014; Hör et al., 2020; Smirnov et al., 2016)**.

Another identified RBP candidate for sRNA-mRNA regulation is the Carbon Storage Regulatory A protein (CsrA). CsrA is the major protein regulator of the Carbon Storage Regulatory (Csr) Network, also known as the Rsm (<u>R</u>egulator of <u>S</u>econdary <u>M</u>etabolites) Network. Overall, the Csr network represses stationary phase processes, including biofilm formation and glycogen synthesis, while activating exponential phase processes such glycolysis and motility (**Romeo, T., & Babitzke, P. 2018**). CsrA is a global RBP widely conserved in the *Gammaproteobacteria* class as well as in the *Firmicutes* and *Planctomycetes* phyla (**Finn et al., 2014, Valkulskas et al., 2015**). CsrA predominantly acts by repressing translation of mRNAs (Figure 1B) by directly occluding the ribosome binding site via binding of a conserved “A(N)GGA” motif contained within a hairpin structure; however, additional cases of CsrA binding and activation of mRNA targets are also documented **(Ren et al. 2014, Renda et al. 2020).** Molecularly, the GGA triplet sequence has been identified as the most critical feature for CsrA binding (**Dubey et al., 2005**); from here we will refer to this as the “GGA Motif”. In *E. coli*, expression of two sRNAs, CsrB and CsrC, antagonize CsrA regulation through binding and sequestration, resulting in the titration of intracellular CsrA concentrations **(Romeo et al., 2013)**. The CsrB and CsrC “sponge” sRNAs have multiple copies of the hairpin contained GGA motif and bind CsrA in stoichiometric ratios up to 9:1 and 5:1, respectively (**M. Y. Liu et al., 1997; Weilbacher et al., 2003**). One additional sRNA known to sequester CsrA is McaS, albeit to a lesser extent than CsrB and CsrC (Figure 1C) (**Kavita et al., 2018**). McaS contains four copies of the GGA CsrA binding motif and can bind two copies of CsrA *in vitro.* Additionally, it co-purifies with CsrA *in vivo* (**Jorgensen et al., 2013**). So far, this sequestration and titration of CsrA has been the main function attributed to CsrA-sRNA interactions in the literature.

In recent years, high-throughput characterizations of the Csr system in *Salmonella* and *E. coli,* identified several additional sRNAs that could directly interact with the CsrA protein (Figure 1C) (**Holmqvist et al., 2016; Potts et al., 2017; Sowa et al., 2017**). Additionally, during the preparation of this manuscript the sRNA Spot42 was identified to have CsrA-dependent interactions, in which CsrA occludes an RNAse E degradation site to prevent degradation (**Lai et al., 2022**). The implication of CsrA-Spot42 interactions is particularly interesting due to the many roles of Spot42 sRNA in cell regulation (**Beisel et al., 2011; Aoyama et al., 2022**). Lastly, work in *Bacillus subtilis* also supports that CsrA-sRNA interactions may play a direct role in impacting sRNA-mRNA regulation (**Müller et al., 2019**). Given documented sRNA association (**Valkulskas et al., 2015**) and (ii) broad conservation (**Romeo, T., & Babitzke, P. 2018**), it has been proposed that CsrA could fulfill a broader regulatory role within sRNA-regulated networks, particularly in organisms lacking Hfq and ProQ, such as *Bacillus subtills* (**Müller et al., 2019**).

In this work, we investigate potential direct interactions and regulatory dependence of sixteen sRNAs with CsrA in *E. coli*; these sRNAs were originally identified by computational prediction studies (SI Table 3) and CLIP-seq studies **(Potts et al. 2017)**. Of the initial 16 sRNAs, we demonstrate 10 novel sRNA-CsrA binding interactions *in vitro,* interestingly some of which do not contain the canonical GGA motif. From this subset, we systematically evaluate the ability of these sRNAs to affect CsrA-mRNA regulation *in vivo* by utilizing overexpression fluorescent reports assays. We also investigate the ability of CsrA to impact the regulatory roles of the sRNAs on their respective target mRNAs. Two sRNAs (FnrS and SgrS) were discovered to serve as “sponges” and sequester CsrA, albeit with reduced levels of “sponging activity” relative to that of the CsrB and CsrC sRNAs from *in vivo* reporter-based assays. Additionally, we found that the presence of CsrA uniquely impacts regulatory activity of three sRNAs (McaS, MicL and Spot42) on their respective mRNA targets. For instance, CsrA enhances McaS-*csgD* and is required for Spot42-*fucP* repression. As a whole, our work expands the post-transcriptional regulatory roles of CsrA beyond its traditional (mRNA-focused) modes of regulation.

## Results

### Identifying additional sRNAs implicated in CsrA regulatory interactions

We compiled a list of 16 sRNAs (Table 1), outside of CsrB, CsrC, and McaS, from previous studies shown to be enriched in a CsrA CLIP-Seq (**Potts el al., 2017**), as well as differentially expressed in a *ΔcsrBΔcsrC* strain compared to wild type *E. coli* (**Sowa et al., 2017;** and re-analyzed here described in Materials and Methods). The set of potential CsrA-binding sRNAs in Table 1 encompasses the diverse activity of known sRNAs in *E. coli*. Eleven of the 16 sRNAs are trans-acting sRNAs that have at least one confirmed mRNA target (Table 1, Supplementary Table S1) and many are associated with specific stress responses or metabolic pathways (Supplementary Table S2). As a note, three of these sRNAs do not contain a GGA binding motif and may be interacting through degenerate GGN-like sequences. To identify putative binding sites, we applied a previously developed biophysical model of CsrA-RNA interaction (**Leistra et al. 2018**). A full computational workflow for using the model is in the Materials and Methods. While some of these sRNAs have been tested for CsrA binding *in vitro* (summarized in Table 1), potential *in vivo* regulatory interactions with CsrA have not been considered, excluding McaS, CsrB, and CsrC. McaS and CsrB are included as positive controls because of their previously established sponge activity for CsrA (**Jorgensen et al., 2013; Dubey et al., 2005**). McaS is considered especially relevant as it contains 4 copies of the GGA motif which is on the same order of magnitude as the other sRNAs of interest; this is in contrast to CsrB, which contains 22 copies of the hairpin-contained GGA motif and is therefore assumed to bind CsrA with higher affinity.

**Table 1.**
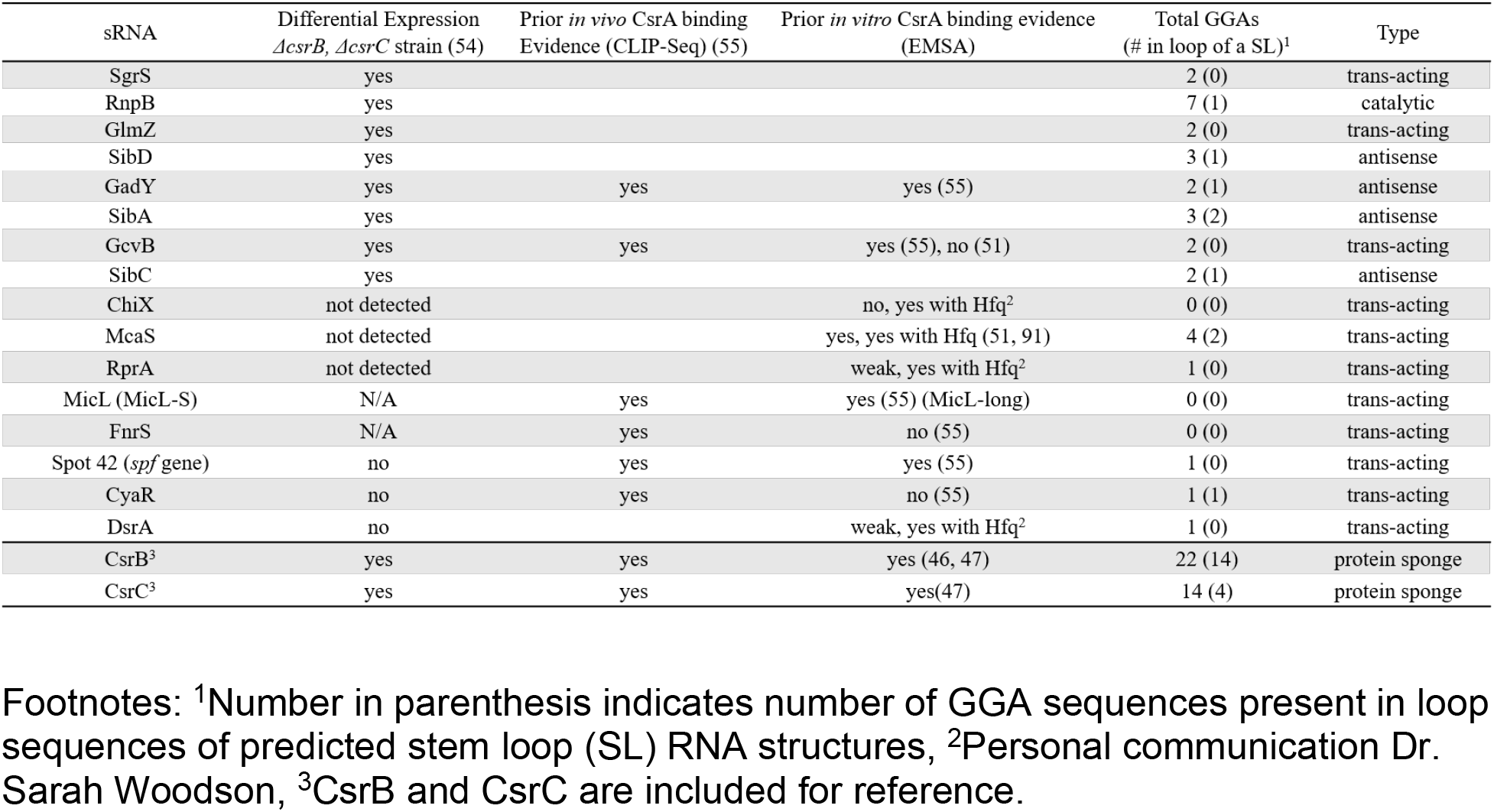
A summary of sRNAs implicated in the Csr system.

However, it is important to note, that the sequence and structural features of the motif are not strict requirements for CsrA binding as degenerate GGA-like sequences may contribute to CsrA-RNA binding and regulation in some target mRNAs. For instance, CsrA binds at non-stem loop GGA sequences in *cstA in vitro* (**Dubey et al., 2003**) and degenerate GGA-like sequences are thought to contribute to CsrA-RNA binding and regulation of some targets (**Kulkarni et al., 2014; Leistra, Gelderman, et al., 2018; Mercante et al., 2009; X. Wang et al., 2005; H. Yakhnin, Yakhnin, et al., 2011**).

### Ten novel sRNA binding partners are confirmed for CsrA *in vitro*

We first assessed whether the 16 identified sRNAs bound CsrA *in vitro* using electrophoretic mobility shift assays (EMSA). CsrA was purified as described in the Materials and Methods and purity was assessed by SDS-PAGE (Supplementary Figure S1). For these experiments, 10 nM of P^32^ – radiolabeled sRNAs were incubated with 3 µM or 6 µM of CsrA, which produced a CsrA:sRNA molar ratio of 300:1 and 600:1, respectively. These ratios were selected based on the reported binding affinities for Spot42, GadY, GcvB, and MicL presumed to be physiologically relevant via CsrA CLIP-Seq data (**Potts et al., 2017**). Additionally, all binding assays are performed in large excess of yeast total RNA, which has been previously shown to inhibit non-specific association of CsrA to labeled sRNA (**A. V Yakhnin et al., 2012**).

Across all conditions, super-shifted complexes indicating CsrA binding were detected upon addition of CsrA for 14 of 16 sRNAs (Figure 2 and Supplementary Figure S2). Clear *in vitro* CsrA binding not yet been reported in the literature was detected for several sRNAs: SgrS, ChiX, RprA, DsrA, and SibA (Figure 2A, B, D, E and Supplementary Figure S2D). In addition to these five novel interactions, the sRNAs Spot 42, GadY, and McaS bound CsrA (Figure 2G, H and I) as has been previously shown, which gives confidence to our EMSA conditions generating true novel sRNA-CsrA interactions. We also observed CsrA binding to MicL, RnpB, GlmZ and SibD at the 6 µM concentration of CsrA, which we believe to be sufficient evidence of *in vitro* CsrA interaction (Supplementary Figure S2A, B, C and E). It is worth noting that stronger *in vitro* CsrA binding has been previously detected for MicL (**Potts et al., 2017**); however, this prior study utilized the full-length sequence (MicL-long), while we tested the short, 5’-processed MicL-S sequence (referred to here as MicL for simplicity). The latter is thought to be the functionally active form of the sRNA (**Guo et al., 2014**). In contrast, CsrA binding was not detected for the GcvB and SibC sRNAs (Supplementary Figure S2F and G). Full EMSA images are included in Supplementary Figure S3.

**Figure 2.**
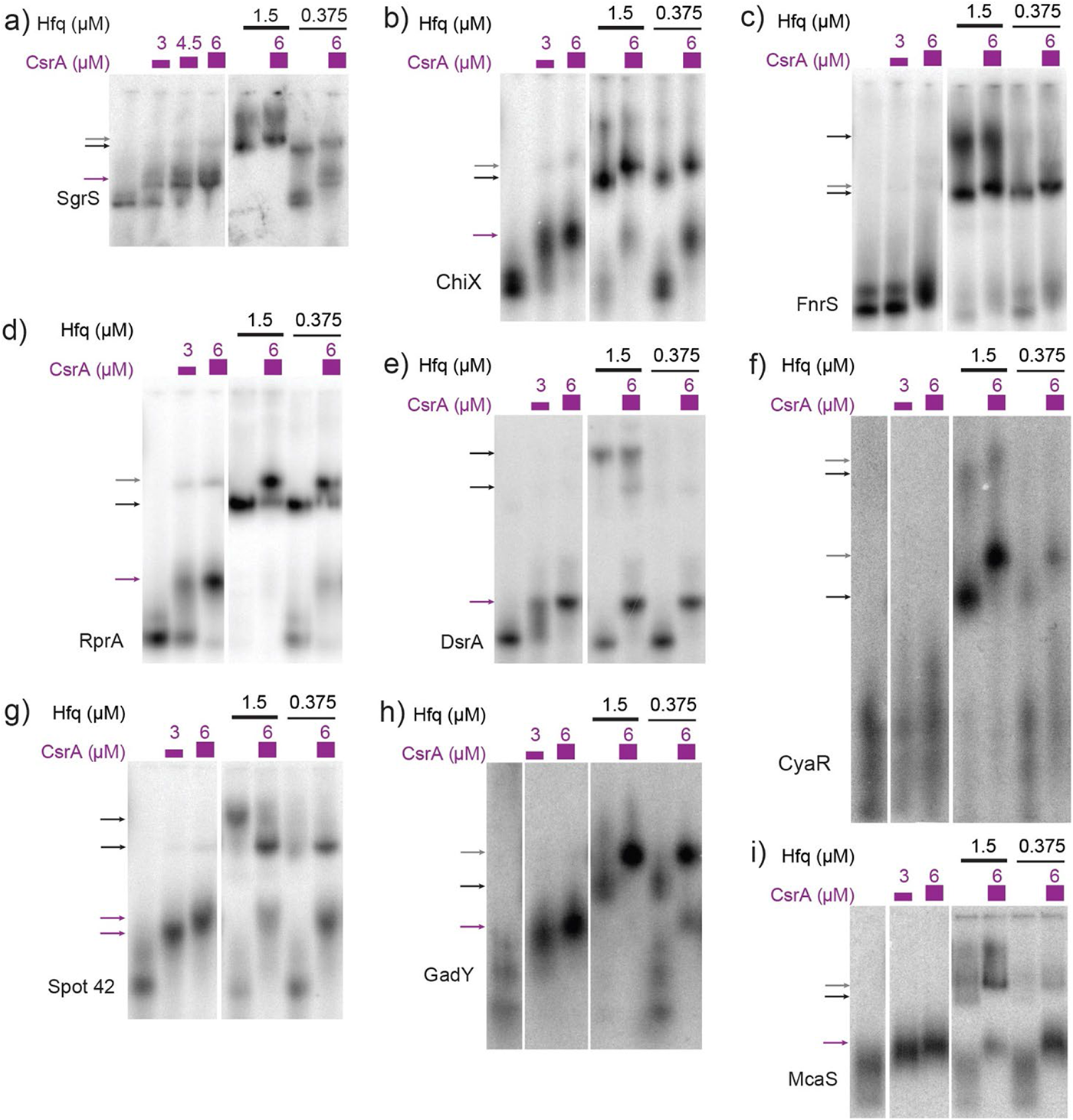
sRNAs bind CsrA in vitro. *In vitro* evaluation of 14 sRNAs with CsrA using Electrophoretic Mobility Shift Assays (EMSAs). These assays were performed by titrating CsrA concentrations (purple), as well as in the presence and absence of Hfq (black). The sRNAs tested are as follows: 2a) SgrS, 2b) ChiX, 2c) FnrS, 2d) RprA, 2e) DsrA, 2f) CyaR, 2g) Spot42, 2h) Gady, 2i) McaS.

Recently, co-immunoprecipitation study of Hfq in *E. coli* demonstrated that CsrA co-precipitates with Hfq in an RNA-dependent manner (**Caillet et al., 2019**). To qualitatively assess if sRNAs from this set can bind both CsrA and Hfq *in vitro* and/or if Hfq influenced CsrA-sRNA binding, we tested binding of all the 16 sRNAs with CsrA (6 µM) in the presence of different Hfq concentrations (0.375 µM or 1.5 µM). These concentrations were selected to mimic estimated ratios natively occurring between these two proteins *in vivo* (Materials and Methods). Importantly, we found that the presence of Hfq affected the binding of many sRNAs to CsrA, likely forming ternary complexes of CsrA-sRNA-Hfq (Figure 2A-E, H and I). These results agree with prior *in vitro* observation of ternary CsrA-sRNA-Hfq complexes for ChiX, RprA, and McaS (**Jorgensen et al., 2013; Peng, 2014 and Dr. S Woodson, personal communication, August 2018**). The complete summary of the EMSA results is compiled in Table 2. In total, we demonstrate 10 novel, and 14 total sRNA-CsrA interactions *in vitro*.

**Table 2.**
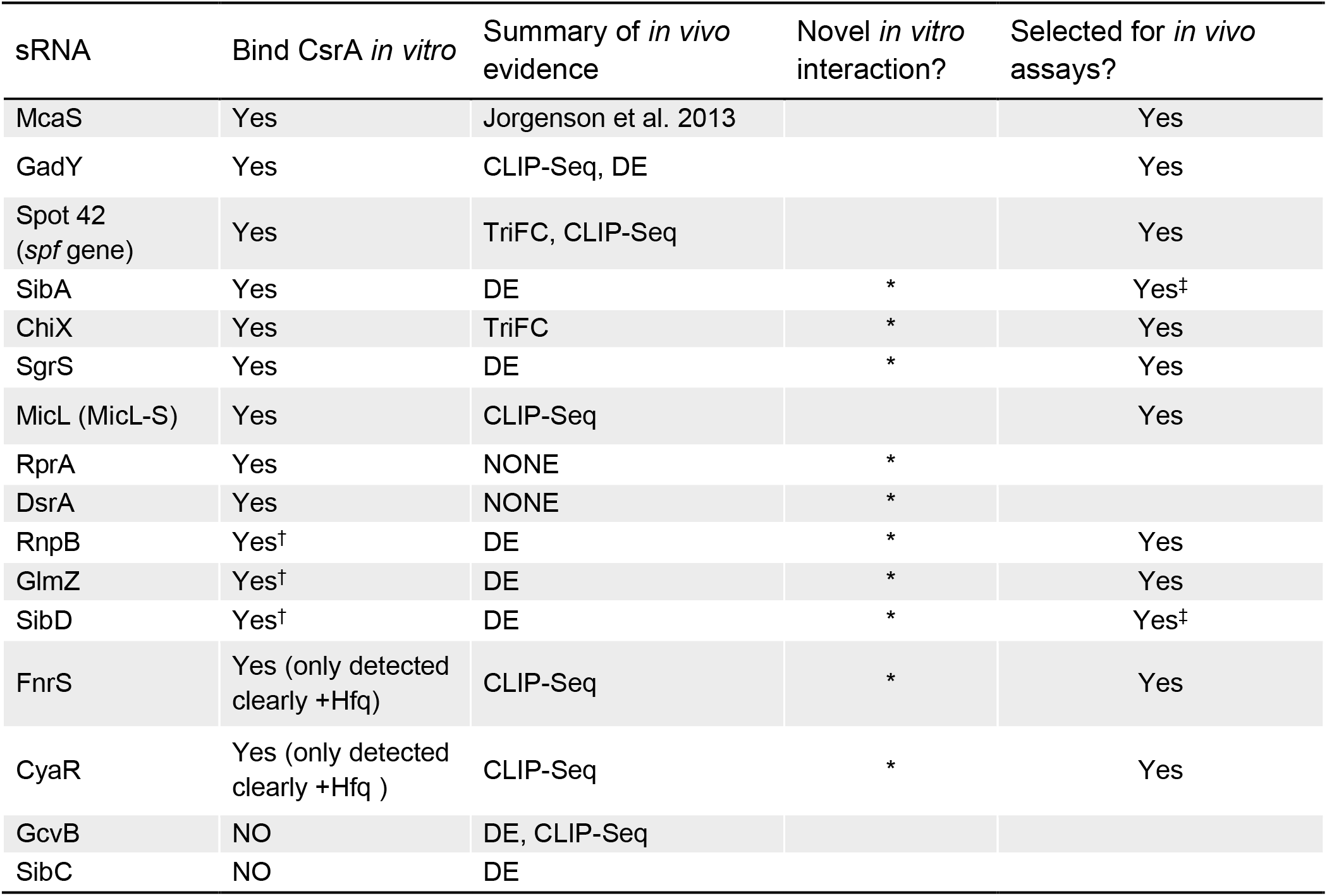
Summarized in vitro sRNA-CsrA binding results from EMSAs.

| sRNA | Bind CsrA *in vitro* | Summary of *in vivo* evidence | Novel *in vitro* interaction? | Selected for *in vivo* assays? |
| --- | --- | --- | --- | --- |
| McaS | Yes | Jorgenson et al. 2013 |  | Yes |
| GadY | Yes | CLIP-Seq, DE |  | Yes |
| Spot 42 (*spf* gene) | Yes | TriFC, CLIP-Seq |  | Yes |
| SibA | Yes | DE | * | Yes<sup>‡</sup> |
| ChiX | Yes | TriFC | * | Yes |
| SgrS | Yes | DE | * | Yes |
| MicL (MicL-S) | Yes | CLIP-Seq |  | Yes |
| RprA | Yes | NONE | * |  |
| DsrA | Yes | NONE | * |  |
| RnpB | Yes<sup>†</sup> | DE | * | Yes |
| GlmZ | Yes<sup>†</sup> | DE | * | Yes |
| SibD | Yes<sup>†</sup> | DE | * | Yes<sup>‡</sup> |
| FnrS | Yes (only detected clearly +Hfq) | CLIP-Seq | * | Yes |
| CyaR | Yes (only detected clearly +Hfq ) | CLIP-Seq | * | Yes |
| GcvB | NO | DE, CLIP-Seq |  |  |
| SibC | NO | DE |  |  |

### *In vivo* sRNA screen identifies CsrA-sequestration activity for FnrS and SgrS

We next investigated the *in vivo* functional implications of the *in vitro* confirmed CsrA-sRNA interactions. We consider two non-exclusive hypotheses: (i) the possibility that sRNAs are minor antagonists of CsrA similar to CsrB and CsrC and (ii) the possibility that sRNA-CsrA binding regulates downstream sRNA-mRNA target interactions. As such, we initially pursued 12 of the 14 sRNAs that (i) bound CsrA *in vitro* and (ii) had some evidence of CsrA impact *in vivo* (Summarized in Table 2). We excluded SibA and SibD as these sRNAs proved difficult to work with *in vivo* due to toxic peptide function (see Materials and Methods). This ultimately left us with 10 total sRNAs to pursue in these assays: McaS, GadY, Spot42, ChiX, SgrS, MicL, RnpB, GlmZ, FnrS, and CyaR.

To evaluate if the found *in vitro* binding interactions could be directly recapitulated *in vivo*, we employed a plasmid-based fluorescence complementation system previously developed in our lab and previously shown to capture CsrA-CsrB interactions (**Gelderman et al., 2015; Leistra, Mihailovic, et al., 2018; Sowa et al., 2017**). Under the conditions tested, we were only able to detect a significant increase in GFP complementation (indicative of sRNA-CsrA proximity) for Spot42-CsrA and ChiX-CsrA, relative to the case of CsrA and a random sRNA sequence control (Supplementary Figure S4B). While allowing us to validate the new *in vivo* interaction between ChiX and CsrA, the YFP complementation signal was low even for the McaS positive control. This may be due to the large MS2 binding protein affecting RNA folding. Thus, these results suggest the need for a more sensitive *in vivo* approach to investigate the *in vitro* supported sRNA-CsrA interactions.

As observed previously for CsrB, CsrC, and McaS (**Jorgensen et al., 2013; M. Y. Liu et al., 1997; Weilbacher et al., 2003**), we anticipated that some of the novel sRNA-CsrA interactions would fall into our first hypothesis and act as a “sponge” sRNA and sequester CsrA *in vivo*. The sponging would then impact regulation of CsrA on its mRNA targets. To screen for this “sponge” activity of each sRNA, we adapted a plasmid-base screen previously utilized in our lab (**Sowa et al., 2017**, **Leistra et al., 2018**). In short, we induced expression of each sRNA with a constitutively expressed *glgC-gfp* translational fusion reporter both K-12 MG1655 wild type *E. coli* and a Δ*csrBCD csrA::kan* strain (*i.e*. Csr system deletion strain). The *glgC-gfp* fusion was selected as CsrA binds and represses translation of the *glgC* mRNA and *glgC* is one of the most well-characterized mRNA targets of CsrA (**Baker et al., 2002; Mercante et al., 2009; Sowa et al., 2017**). As a negative control, a random 80 nt sequence absent of any GGA motifs was selected from the *E. coli* genome (Supplementary Table S1S4). In this system, if a specific sRNA sequesters CsrA (diluting its effective concentration), we would expect to observe an increase in fluorescence (indicative of less repressed *glgC-gfp* expression) following sRNA induction, relative to the fluorescent output of the uninduced system.

Screening revealed that CsrB, McaS, SgrS, and FnrS all significantly increased fluorescence of the *glgC*-*gfp* translational fusion only in the presence of induced CsrA (Figure 3D and E, P-value < 0.05). While CsrB and McaS are positive controls and were expected, SgrS and FnrS are novel findings. Induction of CyaR and ChiX increased *glgC-gfp* fluorescence in the presence of CsrA (Figure 3D, P-value < 0.05 by paired t-test), they also do so in the absence of CsrA (Figure 3E, P-value < 0.05 and P-value =0.06), suggesting that these sRNAs may regulate the translational reporter directly or indirectly via a non-CsrA factor. Surprisingly, GadY does not exhibit sponge activity for CsrA in our assays, as previously reported **(Parker et al., 2017b)**. This discrepancy may be explained by the lower copy number, and thus greater sensitivity, of the CsrA target reporter in the previous work.

**Figure 3.**
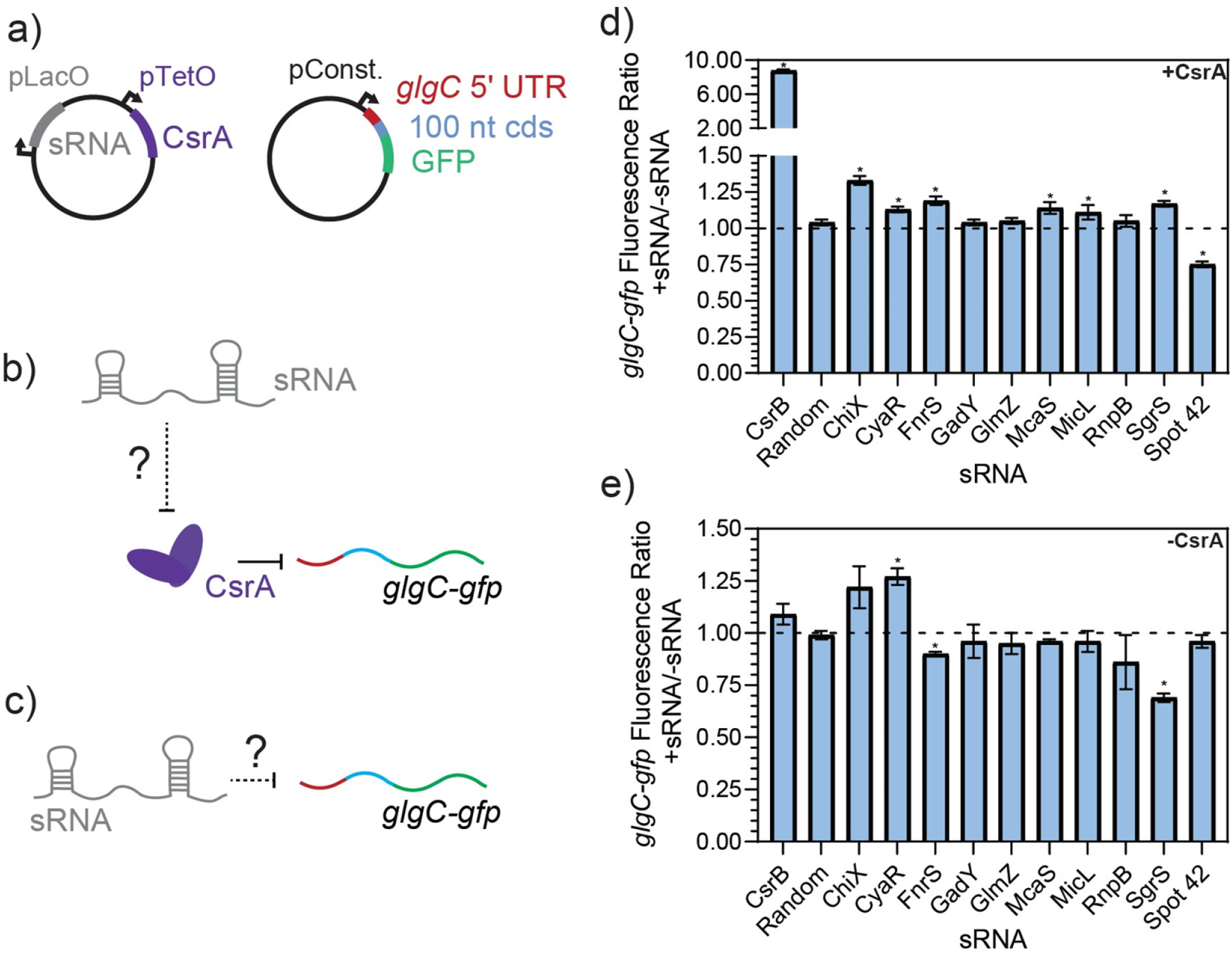
FnrS, SgrS, McaS, and Spot42 alter fluorescence of *glgC-gfp* in a CsrA-dependent manner. 3a) Plasmid schematics of *in vivo* reporter assays to evaluate the effects of an sRNA on CsrA regulation of a known target, the 5’ UTR of the *glgC* mRNA. The 5’ UTR of the *glgC +* 100 nts of the CDS were fused to *gfp* and constitutively expressed from a plasmid. CsrA and each sRNA were expressed from a second plasmid under aTc and IPTG-inducible control, respectively. 3b) Each sRNA may interact with CsrA and affect CsrA repression of the *glgC-gfp* mRNA fusion. 3c) Additionally, each sRNA may interact with the mRNA fusion directly. 3d) Fluorescence ratio of the *glgC-gfp* mRNA fusion between the presence and absence of each sRNA, as well as the presence of CsrA. 3e) Fluorescence ratio of the *glgC-gfp* mRNA fusion between the presence and absence of each sRNA, in the absence of CsrA.

To further evaluate the hypothesis that SgrS and FnrS are two novel CsrA sponge sRNAs, we interrupted putative CsrA binding sites within these sRNAs expecting this would abrogate CsrA sponge activity. McaS was used as a positive control in these experiments, given its earlier (i) demonstration of CsrA sponge activity, (ii) confirmation of CsrA binding sites, and (iii) more similar number of GGA motifs (compared to CsrB). Likewise, CyaR was used as a negative control because we anticipated that disrupting putative CsrA binding sites would not impact the likely CsrA-independent observed increase in *glgC-gfp* fluorescence (Figure 3D and E). For the sRNAs that contain one or more instances of the conserved GGA binding motif, we performed GGA:CCA mutations as done in previous literature to interrupt CsrA-RNA binding (**Dubey et al., 2005; Patterson-Fortin et al., 2013**). It should be noted that alternative mutations were made to the GGA motif if the standard GGA:CCA mutation altered the predicted secondary structure of the sRNA (Materials and Methods). We disrupted each instance of the GGA motif in the SgrS and CyaR sRNAs (termed “GGA mutants”, Supplementary Table S4). For McaS, the two GGA motifs (of 4 total) that were shown to be CsrA binding sites *in vitro* and *in vivo* in a previous study (**Jorgensen et al., 2013**) were mutated (“dual GGA mutant,” Supplementary Table S1S4). For FnrS, which does not contain a GGA motif, GGN sequences within the most likely pair of putative CsrA binding sites predicted using the biophysical model (Supplementary Table S3) were mutated (“GGN mutant 1,” Supplementary Table S4).

The mutant sRNAs were screened in the same plasmid assay. Induction of mutant FnrS, SgrS, and McaS mutant sRNAs significantly reduced fold increase of *glgC-gfp* fluorescence relative to the corresponding wild type sRNA (Figure 4A). Northern blotting analysis of these samples indicated both the wild type and mutant sRNAs are induced at comparable levels (Figure 4B, C and D). These data agree with a previous study of McaS, where mutations of the same GGA motifs significantly decreased its CsrA sponge activity, as measured by a *pgaA-lacZ* reporter (**Jorgensen et al., 2013**). Additionally, as expected, the CyaR mutant sRNA does not alter *glgC*-*gfp* fluorescence between the wild type and mutant sRNAs (Figure 4A), supporting the hypothesis that the observed effects of CyaR on *glgC*-*gfp* fluorescence (Figure 3D and E) do not involve direct CyaR-CsrA interactions. Lastly, altering the known GGA motif in Spot42 and expressing the mutant sRNA slightly alleviates the fold-repression effect observed when using the wild type Spot42 sRNA sequence, though that difference is not statistically significant (Figure 4A). This does point to the idea that the decrease in *glgC-gfp* fluorescence upon expression of Spot42 relies upon a direct CsrA-sRNA interaction.

**Figure 4.**
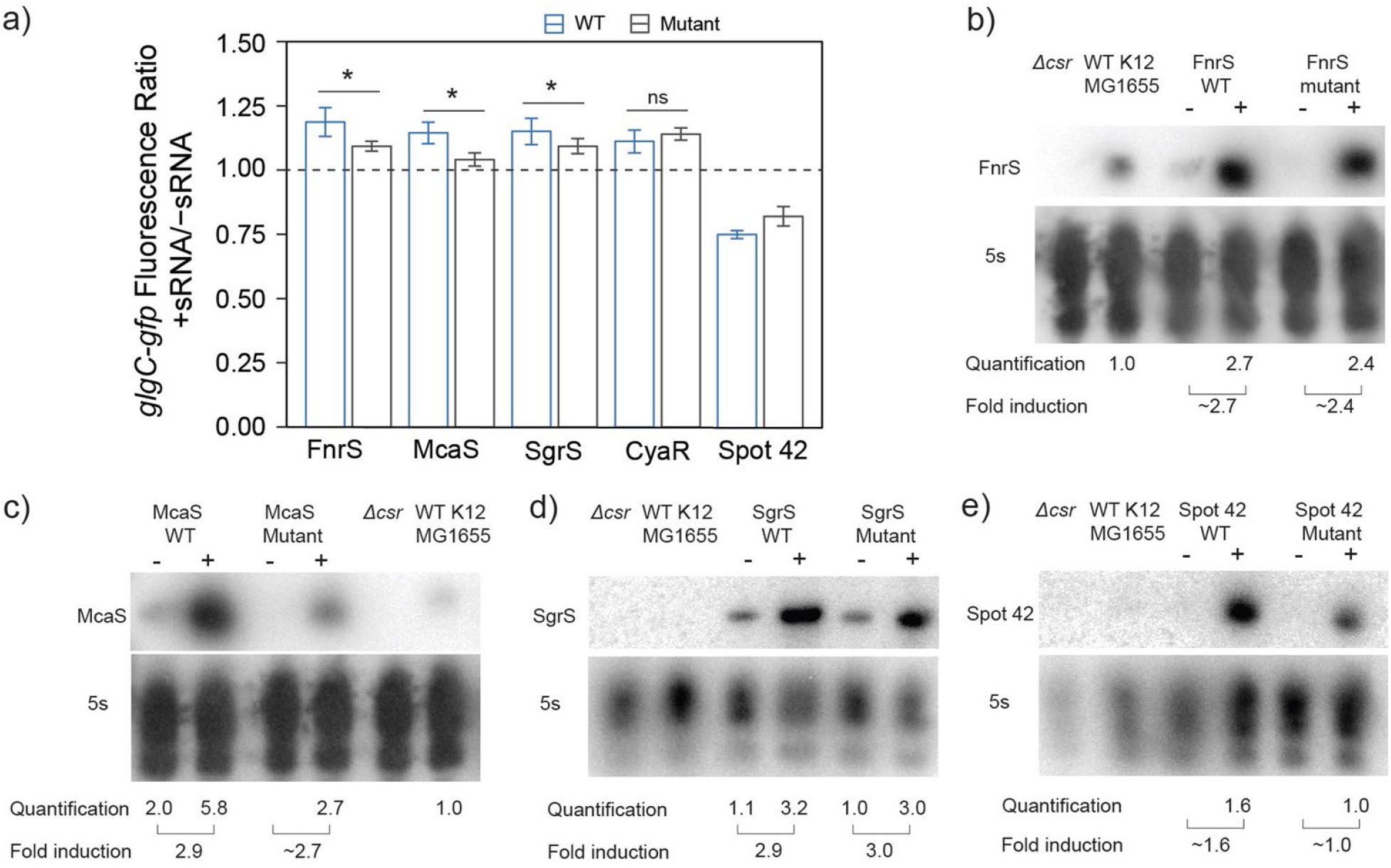
*In vivo* mutational assays confirm that direct sRNA-CsrA interaction enables CsrA-sponging activity of McaS, SgrS, and FnrS. 4a) Fluorescence ratio of the *glgC-gfp* mRNA fusion reporter between the presence and absence of the FnrS, McaS, SgrS, CyaR, and Spot42 sRNAs. Both the wild type (blue) and mutant (grey) sRNAs were tested. For descriptions of how the sRNAs mutants were generated to abrogate CsrA interactions see Materials and Methods. 4b-e) Northern Blots of each wild and mutant sRNA to ensure consistent stability between the wild type and mutant sRNAs.

The above data indicate that SgrS and FnrS can exert CsrA sponge activity similar to that of McaS in a plasmid-based system. However, the effects of all of these sRNAs on *glgC-gfp* fluorescence are relatively small (∼1.2-fold increases) (Figure 4A) compared to that observed for CsrB (∼9-fold increase) (Figure 3D). This drastic difference is expected, as CsrB contains 22 GGA CsrA binding motifs while McaS, Sgrs, and FnrS contain only 4 CsrA binding motifs at most (Summarized in Table 1). It should also be noted that, when assessing the impact of SgrS and McaS on endogenous CsrA regulation of the *glgC* and *pgaA* mRNAs by qRT-PCR, we observed no significant impact on *glgC* or *pgaA* abundance for McaS nor SgrS (Supplementary Figure S6). However, these experiments were conducted under common laboratory growth conditions (mid-exponential phase in LB media). Previous study of McaS demonstrated greater expression of McaS than CsrB and CsrC in late stationary phase growth in colonization factor antigen media and hypothesized that McaS-CsrA sequestration may be relevant under this growth condition (**Jorgensen et al., 2013**). This observation raises the question of which physiological conditions these additional “sponges” exert biologically relevant sequestration activity on CsrA to affect regulation of its mRNA targets.

### CsrA impacts regulatory activity of the McaS sRNA on its mRNA targets

We next evaluated our second hypothesis: that CsrA remodels the downstream regulatory interactions of the sRNA regulatory network itself. While this work was in preparation, CsrA enhancing Spot 42-*srlA* mRNA repression was elucidated (**Lai et al., 2022**), setting the precedent that CsrA can remodel downstream sRNA networks. In light of these possibilities, we tested the impact of CsrA on sRNA-mRNA regulation for CyaR-*yobF*, MicL-*lpp*, Spot 42-*fucP*, Spot 42-*ascF*, SgrS-*manX*, SgrS-*ptsG*, FnrS-*folX*, and McaS-*csgD*. This subset included all sRNAs with multiple confirmed mRNA targets that we identified to interact with CsrA *in vitro* and had prior evidence of affecting CsrA *in vivo* (Table 2). It should be noted that we attempted to select the mRNA targets most sensitive to each sRNA regulator *in vivo* according to literature, if it was characterized (e.g., *ptsG* for SgrS **(Bobrovskyy et al., 2019)**). We excluded testing ChiX as we were unsuccessful at constructing reporters for the ChiX-mRNAs *citA* and *chiP* to due to the fact the translational fusions were not fluorescent.

To evaluate the effect of CsrA on sRNA-mRNA regulation, we adapted the fluorescence assay described previously (Figure 3A) to determine the impact of sRNA-CsrA interaction on sRNA-mRNA regulation. Briefly, the *glgC* 5’ UTR was replaced with the 5’ UTR of at least one confirmed mRNA target for each sRNA and constitutively expressed in the Δ*csrBCD csrA::kan* strain. Both the sRNA and CsrA were induced on plasmids individually and concurrently to determine the impact of CsrA on sRNA-mRNA regulation.

After confirming that expected sRNA-mRNA repression was detectable for all pairs except CyaR-*yobF* (Supplementary Figure S8), we noticed that in some cases, the presence of CsrA affected the sRNA’s ability to regulate its cognate mRNA target. We first observed that McaS-*csgD* repression is significantly enhanced in the presence of CsrA. While it has been shown that McaS sponges CsrA to affect expression of CsrA targets in a CsrA-dependent manner (e.g., *pgaA*) (**Jorgensen et al., 2013**), there has been very little work into understanding if CsrA affects the metabolic networks that McaS itself regulates. McaS has been shown to target the *csgD* and *flhD* mRNAs, which are master transcriptional activators of curli biogenesis and flagella synthesis (**Thomason et al., 2012**). From our reporter assay we observed *csgD*-*gfp* fluorescence is significantly lower in the presence of both McaS and CsrA, relative to McaS alone (Figure 5A, T-test P-value < 0.05). Although *csgD-gfp* fluorescence increases upon induction of CsrA alone, this change is likely an indirect effect of expressing the global CsrA regulator. When each condition is compared to its respective CsrA null condition (+sRNA –CsrA to –sRNA –CsrA; +sRNA +CsrA to -sRNA +CsrA), repression of the *csgD* reporter significantly enhanced upon expression of McaS alongside CsrA (Figure 5A, T-test P-value < 0.05).

**Figure 5.**
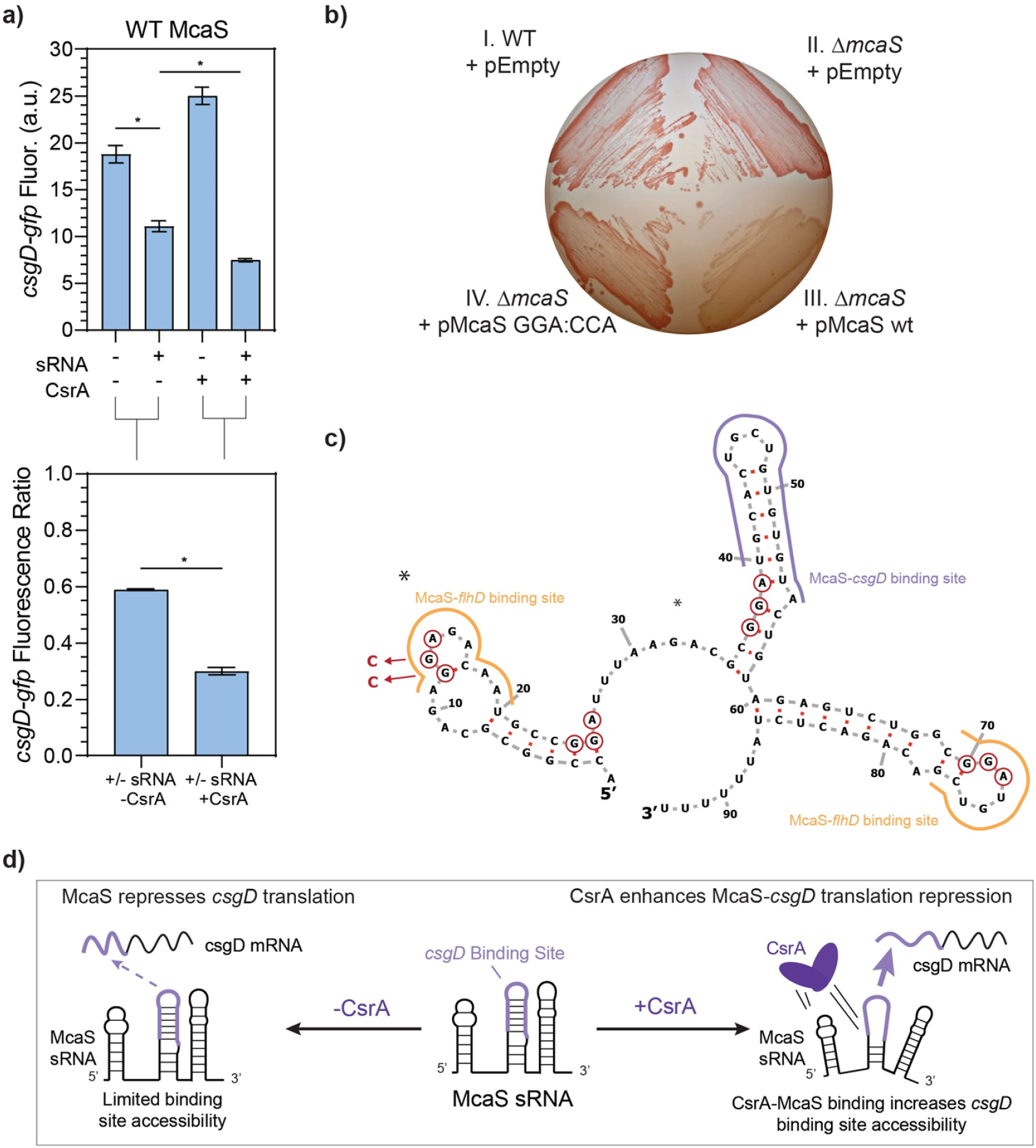
McaS-*csgD* repression is enhanced by direct CsrA interactions under native expression conditions. 5a) Fluorescence of a *csgD-gfp* mRNA reporter in response to both the McaS sRNA and CsrA expression tested combinatorially. The fluorescence ratios of the *csgD-gfp* reporter between the presence and absence of McaS are calculated with and without CsrA expression. The *csgD-gfp* mRNA reporter was constructed similarly to the *glgC-gfp* mRNA reporter. 5b) Effect of CsrA on McaS-csgD regulation evaluated by the Congo Red plate to measure endogenous curli expression. (Quadrant I) Wild type *E. coli* transformed with -an empty control version of a low expression plasmid (pEmpty), (II) *ΔmcaS E. coli* transformed with pEmpty, (III) pMcaS wild type, and (IV) pMcaS mutant (5’-most GGA:CCA) were grown for 48 hours with limited NaCl as to induce native curli expression. Plasmid-based induction of wild type McaS (1mM IPTG) effectively repressed the *csgD* mRNA (white colonies) compared to wild type *E. coli* (dark red colonies), while mutant McaS (5’-most GGA:CCA) showed minor repression (light red colonies). 5c) Predicted secondary structure of McaS (Vienna RNA), with GGA motifs (red-circled nts), mutations (blue arrows and nucleotides), and known *csgD* and *flhD* mRNA binding sites indicated (purple and yellow outlines, respectively). 5d) Proposed effect of CsrA on McaS-*csgD* regulation: CsrA enhances repression by increasing mRNA-sRNA binding site accessibility.

To assess if direct interaction between CsrA and McaS was leading to the measured repression, a mutant Mcas sRNA was constructed with the 5’-most GGA CsrA binding site mutated (Supplementary Table S4). Using the same assay, Enhanced repression in the presence of CsrA is significantly reduced when using the McaS mutant sRNA (Supplementary Figure S10A). While alleviation of repression is not fully achieved using the McaS mutant, this may be due to the fact that the second known GGA binding site for CsrA in McaS (**Jorgensen et al., 2013**) overlaps with the *csgD* binding site (Figure 5C) and thus not mutated. Importantly, these results suggest that CsrA is in fact remodeling the interaction between McaS and *csgD* via direct McaS-CsrA binding. This is the first time CsrA has been observed to remodel the network of one of its sponge sRNAs. Differential McaS sRNA accumulation between the presence and absence of CsrA was confirmed to be marginal via Northern blotting analysis (Supplementary Figure S10B).

Enhanced McaS-*csgD* repression at physiologically relevant conditions was additionally observed by assaying curli production, a phenotype governed by *csgD* expression, on Congo Red indicator plates. In accordance with previous work (**Thomason et al., 2012**), endogenous expression of *csgD* produces bright red colonies in both wild type and *ΔmcaS E. coli* strains, as endogenous McaS is likely not expressed under the curli-producing conditions (Figure 5B, Panels I and II). Expression of wild type McaS from a low copy number plasmid (Materials and Methods) produces opaque white colonies, indicating *csgD* repression (Figure 5B, Panel III). These opaque white colonies were also observed previously (**Thomason et al., 2012**). Importantly, expression of the McaS mutant (where CsrA interaction is diminished) restores the red colony color observed when *csgD* is not repressed (Figure 5B, Panel IV), suggesting that a functional McaS interaction with endogenous CsrA enhances McaS repression of *csgD* and subsequent curli formation.

We designed this McaS mutant (5’ GGA:CCA, Supplementary Table S4) based on the hypothesis that direct CsrA binding at the 5’-most GGA in McaS is critical to enhanced *csgD* repression. Previous *in vitro* McaS-CsrA binding assays support this notion, as disrupting the 5’-most GGA of McaS impeded CsrA binding more than disrupting the GGA site that overlaps with the *csgD* binding site. Based on these data, we propose CsrA-McaS binding at the 5’-most GGA motif seeds a bridging interaction that disrupts a downstream hairpin structure containing the *csgD* binding site (Figure 5D). Such a McaS-CsrA complex could increase accessibility of the McaS binding site for *csgD* and enhance *csgD* repression. Alternatively, since McaS-CsrA binding at the 5’-most GGA motif directly overlaps a known McaS-*flhD* mRNA binding site (Figure 5E), it is plausible that McaS-CsrA interaction may bias partitioning of McaS among its mRNA targets towards *csgD* rather than *flhD.* We also looked to map the impact of McaS-CsrA binding on the only other known target of McaS, *flhD*. However, a sufficiently fluorescent *flhD*-*gfp* reporter construct could not be constructed.

### CsrA binds Spot42 at two additional non-GGA sites and is required for Spot42 regulation of its mRNA target *fucP* under physiologically relevant conditions

In addition to remodeling McaS-*csgD* repression, we observed that CsrA significantly impacts Spot 42-*fucP* regulation in our plasmid-based assay. Specifically, Spot 42-*fucP* repression was only observed with the *fucP-gfp* reporter when CsrA expression was induced while apparent significant *fucP* activation was detected with the *fucP-gfp* reporter in the absence of CsrA (Figure 6A). Given that for these overexpression reporter assays (Figure 6A), CsrA was expressed on medium copy number plasmids leading to component concentrations 5-15-fold greater than native levels as previously quantified in (**Sowa et al. 2017**), we next evaluated if CsrA-dependent regulation could be observed under more physiologically relevant conditions.

**Figure 6.**
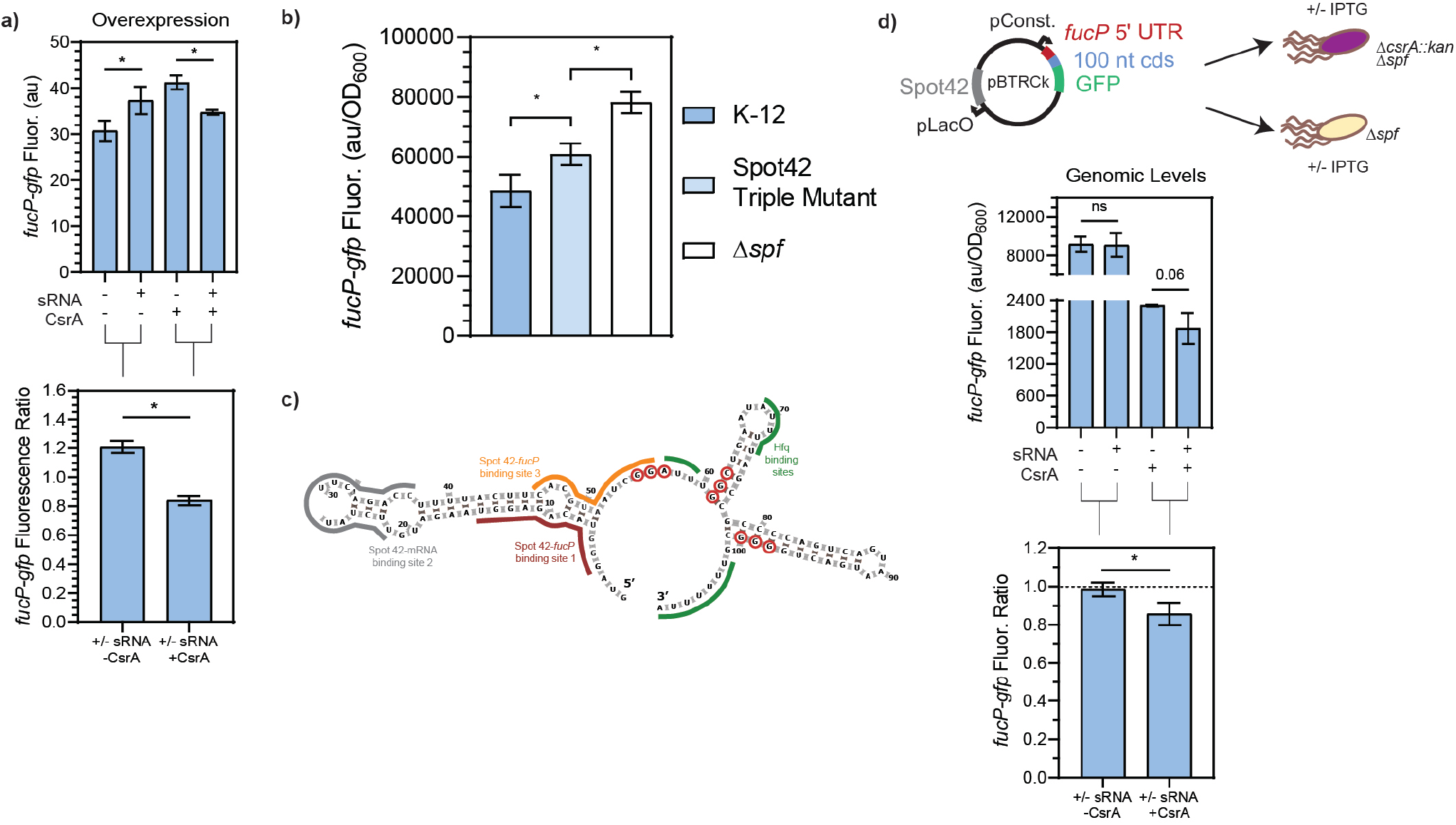
CsrA is necessary for Spot42 regulation of *fucP* target under native conditions. 6a) Fluorescence of a *fucP-gfp* mRNA reporter in response to both the Spot42 sRNA and CsrA expression tested combinatorically. The fluorescence ratios of the *fucP-gfp* reporter between the presence and absence of Spot42 are calculated with and without CsrA expression. The *fucP-gfp* mRNA reporter was constructed similarly to the *glgC-gfp* mRNA reporter. 6b) Expression of the *fucP-gfp* mRNA reporter in WT, a mutant Spot42, and a Δ*spf* strain. The mutations made to Spot42 reduced but did not fully eliminate CsrA-Spot42 interactions. In this assay, cultures containing the *fucP-gfp* reporter plasmid were grown in minimal media and Spot42, when present, was expressed from the genome using a bolus addition of glucose. 6c) Predicted secondary structure of Spot42 (Vienna RNA), with GGA or GGN motifs (red-circled nts) and known *fucP* mRNA (maroon and yellow) and Hfq (green) binding sites indicated. 6d) General workflow of low copy plasmid reporter assay and results. Fluorescence a *fucP-gfp* mRNA reporter in response to Spot42 and CsrA expression in a genomic context. The *fucP-mRNA* reporter was constitutively expressed from a low copy plasmid, while Spot42 is induced from the same plasmid. This was tested in the Δspf and ΔspfΔ*csrA::kan* strains of *E. coli*, to evaluate Spot42-*fucP* regulation under native CsrA expression. Ratios of the *fucP-gfp* reporter we calculated in the presence and absence of CsrA for the WT and *csrA::kan* strains.

As a tool to test biological relevance, we required a Spot42 sRNA mutant with abrogated CsrA interactions. We thus mapped the sites on the Spot42 sRNA that contributed to CsrA binding. We first conducted RNase T1 enzymatic probing of CsrA-Spot42 binding (Supplementary Figure S11A) to identify additional potential CsrA binding sites on the Spot42 sRNA, beyond the previously identified one GGA binding site that overlaps one of the RNAse E cleavage sites (**Lai et al., 2022**). Importantly, Spot 42 only contains one GGA CsrA binding motif, so we anticipated that additional binding sites would correspond to degenerate GGA-like sequences. From our sequencing gel, we identified protection on the 3’ region of the Spot42 sRNA, starting at nucleotide 55, in accordance with the previously identified GGA binding motif (Supplementary Figure S11A). Within the rest of the 3’ end of Spot42, we identified a GGC and GGG motif, at nucleotides 61-63 and 97-99 respectively, that showed protection in the presence of CsrA. With the GGA, GGC, and GGG motifs as guides, we ran EMSAs to evaluate the sites of Spot 42-CsrA interaction. Briefly, mutations were made to eliminate the putative binding motifs without disrupting the predicted secondary structure of the Spot42 sRNA (Supplementary Figure S11D). After testing each of the mutations in combination, we observed an ∼ 1.8-fold reduction in Spot42-CsrA binding only when all three mutations were present in the sRNA (Supplementary Figure S11B and C). As such, we conclude CsrA binds Spot 42 using the established GGA site, as well as the two novel degenerate GGN sites on the 3’ end of Spot42. We anticipate that a preferred interaction between Spot 42 and one binding pocket of a CsrA dimer at the GGA site likely tethers a dimer to the sRNA and allows for an additional contact to be made at a lower affinity GGN site by the other binding pocket. We expect that the single stranded GGG site is more likely bound alongside the GGA site, given its greater accessibility than the hairpin contained GGC site (Supplementary Figure S11D). Such a model has been previously proposed for CsrA-RNA binding (**Mercante et al., 2009**) and likely also explains CsrA repression of the *hfq* transcript, which contains only a single GGA site but presents a CsrA-occluded AGA site in *in vitro* footprinting studies (**Baker et al., 2007; Leistra, Gelderman, et al., 2018**). Moreover, degenerate GGA-like sequences (*e.g.,* AG-rich or GGN sequences) are thought to contribute to CsrA-RNA binding and regulation in some target mRNAs (*i.e.*, *glgC*, *csrA*, and *pgaA*), although these targets also contain at least two instances of the preferred GGA CsrA binding site (**Mercante et al., 2009; X. Wang et al., 2005; H. Yakhnin, Yakhnin, et al., 2011**).

Using this information, we designed a strain with Spot42 containing all three mutations to abrogate Spot42-CsrA interactions *in vivo*, a *Δspf::spf_triple_*GGNs mutant (referred to as Spot42 Triple Mutant hereafter). To directly evaluate CsrA-Spot42 interactions upon the regulatory Spot42*-fucP* activity, we measured regulation of the *fucP-gfp* translational reporter expressed from a low copy number plasmid (Materials and Methods) in wild type *E. coli,* the Spot42 triple mutant strain and a *Δspf* strain. We induced endogenous Spot 42 transcription from the genome at an OD_600_ of 0.2 by addition of 0.2% glucose to M9 media supplemented with 0.4% glycerol and casamino acids, alleviating CRP-Spot 42 transcriptional repression, as shown in previous literature (**Beisel & Storz, 2011**). From this assay, we observed that the Spot42 repression of the *fucP-gfp* reporter is significantly alleviated in the Spot42 Triple Mutant expressing strain relative to the wild type Spot42 strain (Figure 6B). These results point to the fact that Spot42 repression of the *fucP-gfp* is in fact partially? CsrA dependent under physiologically relevant conditions. Interestingly, we could not alleviate the fucP-gfp reporter signal to that of the *Δspf* in the Spot42 Triple Mutant strain. This is likely due to network interactions other than CsrA, such as residual Hfq-driven repression of the *fucP-gfp* target by Spot42 (as seen in literature ref).

Lastly to understand if we could observe the CsrA-dependent regulation of the *fucP-gfp* reporter by Spot42 occurred under physiological conditions, we modified the reporter assay from Figure 6A such that it could be recapitulated at genomically relevant CsrA concentrations. As such, we chose to use a *Δspf* and *ΔcsrA::kanΔspf* strain of *E. coli* to replicate the plus and minus CsrA conditions and expressed Spot42 from the low copy plasmid mentioned earlier. This was chosen, as Spot42 cannot be reliably expressed from the genome in a *ΔcsrA::kan* strain (**Beisel & Storz, 2011**). Using this adapted assay, we observed that repression of the *fucP-gfp* reporter by Spot42 only occurred when CsrA was present. (**Figure 6D**). There was no change in *fucP-gfp* reporter signal in the *ΔcsrA::kan*, regardless of Spot42 expression. With these data in mind, we demonstrate that under physiological conditions, Spot42 repression of the *fucP-gfp* transcript is CsrA-dependent. Additionally, while this work was being prepared a destabilizing effect of Spot42 in a *csrA* deletion strain of *E.* coli was published (**Lai et al., 2022**). This work is the second known demonstration of CsrA-dependent regulation of an mRNA target via Spot42.

## Discussion

In this work, we have characterized an extended overlap between the CsrA global regulatory protein and sRNA networks. Our results indicate that a much larger set of mRNA-binding sRNAs bind CsrA *in vitro* (Figure 2 and Table 2) and that sRNA-mRNA regulation can be impacted by CsrA *in vivo.* Importantly, our data support the notion that CsrA-sRNA interactions result in different post-transcriptional regulatory schemes and have selective impacts in the mRNA target network of the sRNA.

Consistent with the notion of sRNAs acting as “sponges” that sequester CsrA *in vivo*, we have also identified FnrS and SgrS as two novel sRNA that sequester CsrA. While we did not see an effect of McaS and SgrS on the endogenous regulation of the *glgC* and *pgaA* mRNAs under standard laboratory growth conditions. It is plausible that there are different growth conditions in which these sRNAs play a larger role in sequestering and titrating CsrA. Furthermore, the effect of FnrS on endogenous regulation of CsrA targets remains to be assessed, as its regulatory function is understood to be limited to anaerobic conditions (**Durand & Storz, 2010**) and thus, it was excluded from our initial experiments under standard lab conditions.

Beyond the sequestration mechanism, our data support novel influence of the global CsrA regulatory protein on the downstream regulatory roles of sRNAs under physiologically relevant conditions: two examples in this work are the enhancement of McaS-*csgD* repression (Figure 5) and the CsrA-dependent repression of *fucP* by Spot42 (Figure 6), involving new regions of Spot 42-CsrA interactions that include non-canonical degenerate GGA sites, which have not been widely reported. From our *in vivo* reporter assays, we observed that expressing McaS and CsrA simultaneously led to enhanced *csgD* repression, relative to *csgD* repression when only expressing McaS (Figure 5A). Furthermore, we demonstrate this regulatory pattern is consistent under physiologically relevant concentrations and has observable phenotypic outputs by assaying curli production (Figure 5B). While McaS has been previously shown to interact with CsrA (**Jorgenson et al. 2013**), McaS was implicated as a sRNA sponge of CsrA. Our work demonstrates a new role of CsrA in the context of remodeling the metabolic network of McaS, as shown at least by the case of McaS-*csgD* regulation. We can begin to deduce the native context in which CsrA may remodel such sRNA-mRNA interactions by considering known expression conditions and activity for each regulator (Supplementary Table S2). For example, high McaS expression coupled with low CsrA availability (in early and mid-stationary phase), may not yield any observable effects on the mRNA target. But McaS expression under high CsrA availability (in late stationary phase) may demonstrate enhanced McaS-*csgD* repression by CsrA, as these conditions allow McaS to effectively compete for CsrA sequestration.

Beyond the role of CsrA on the regulatory roles of McaS and Spot42, we observed that MicL-*lpp* repression is significantly impeded in the presence of CsrA (Supplementary Figure S9); MicL is another well-characterized sRNAs known to be the key regulatory sRNA in outer membrane protein synthesis and can even affect Sigma E transcription factor activity (**Guo et al. 2014**). Notably, the most likely predicted MicL-CsrA binding region overlaps the known MicL-*lpp* binding site (Supplementary Figure S9E). As such, we evaluated the hypothesis that MicL-CsrA binding directly occludes MicL-*lpp* binding *in vitro* and we observed that the presence of CsrA decreased MicL-*lpp* complex formation approximately two-fold in EMSA analysis (Supplementary Figure S9D). However, given the small effect size *in vivo* and overall limited extent of MicL-*lpp* binding *in vitro*, these observations require further work to confirm direct impact of CsrA on MicL-*lpp* regulation.

In the case of Spot42, the impact of CsrA on mRNA regulation varies. In this work, CsrA is required for Spot42-*fucP* regulation (Figure 6) but does not impact *ascF* repression (Supplementary Figure S8), despite *ascF* being another confirmed target of Spot42 (**Beisel et al. 2012**). While this work was in preparation, CsrA-Spot42 binding was shown to enhance Spot42-*srlA* repression *in vivo* (**Lai et al., 2022**). Our observations that Spot42 requires CsrA to repress *fucP* is the second known example of this form of CsrA-dependent sRNA-mRNA regulation and opens the door towards new ways by which CsrA can remodel the global post-transcriptional landscape, as well as the interplay between CsrA and RNAse E networks.

Previously, Spot42 has been shown to be destabilized in a *csrA* mutant strain *in vivo* (**Lai et al., 2022**). Interestingly, we observed that CsrA affects Spot42 regulation of *fucP* (Figure 6) and *srlA* (**Lai et al. 2022**), but not of its other mRNA target *ascF* (Supplementary Figure S8). Our observations on the requirement of CsrA for the selective repression of some Spot42 targets (*i.e.*, *fucP*) by Spot42 are consistent with a hypothesis in which CsrA affects availability of specific target binding sites within the Spot42 sRNA in addition to its general stability. Further support for this hypothesis stems from the observation that Spot42 has non-overlapping binding sites for *ascF* and *galK* mRNA targets but not for *srlA* and *fucP* (**Beisel et al. 2012**). Collectively, this data suggests that CsrA can affect sRNA-mRNA regulation in a selective fashion. Furthermore, given that *In Vitro* Transcription and Translation (IVTT) assays ruled out the possibility of CsrA directly binding the *fucP* in these interactions (Supplementary Figure S12D and S12E), we propose that the general model for CsrA impacting sRNA-mRNA regulation differs from that of Hfq and ProQ in that a CsrA dimer is not anticipated to bind two different RNA molecules at each of its binding faces and seed annealing. Rather, we presume that a CsrA dimer binds at two sites within a single sRNA to directly occlude sRNA-mRNA binding or alter sRNA-mRNA binding availability at a neighboring region. As such, our results show that CsrA interacts with Spot42 via binding at the previously known GGA site, as well as degenerate GGNs at the 3’ end of the sRNA (Figure 6B, Supplementary Figure S12A). This model is consistent with current understanding of CsrA-mRNA regulatory mechanisms **(Mercante et al., 2009; Romeo & Babitzke, 2018)**.

Interestingly, we observed a bi-directional response of Spot42-*fucP* regulation in the presence or absence of overexpressed CsrA in our initial overexpression *in vivo* reporter assays (Figure 6A). In the absence of CsrA, the *fucP-gfp* reporter was activated when co-expressed with Spot42, but in the presence of CsrA, *fucP-gfp* signal was repressed (Figure 6A). While not initially observed in a genomic context (Figure 6B), upon mutating the canonical Spot42-*fucP* binding site on the *fucP-gfp* reporter, we observed the same CsrA-dependent toggling effect at genomic levels (Supplementary Figure S12B). Moreover, our IVTT assays also showed the bidirectional response upon adding Spot42 sRNA and titration of CsrA into these reactions (Supplementary Figure S12D). These results lay the foundation of the possibility that CsrA could be used as a synthetic bi-directional regulator to construct toggle switches. Previously, post-transcriptional circuits that leverage natively expressed proteins with engineered RNAs are some of the most powerful in synthetic biology (**Na et al., 2013; Noh et al., 2017**).

Our work here uncovered several novel interactions of sRNAs binding both CsrA and Hfq in ternary complexes; such binding has been previously reported for McaS (**Jorgensen et al., 2013**) and observed for ChiX and DsrA (personal communication, Dr. Sarah Woodson’s lab). We report new CsrA-Hfq ternary complexes also with SgrS, FnrS, CyaR, GadY, MicL, RnpB, and GlmZ (Figure 2 and Supplementary Figure S2). Several lines of evidence support the hypothesis that these ternary complexes are RNA-mediated, rather than purely dependent on protein-protein interaction. For instance, the sRNAs Spot42, DsrA, GcvB, and SibD bind Hfq but do not show CsrA-sRNA-Hfq ternary complexes. This contrasts with what would be expected if CsrA and Hfq-associated on a protein-only basis. Additionally, it was recently shown that in *E. coli*, CsrA co-purifies with Hfq in an RNA-dependent manner (**Caillet et al., 2019**). Similarly, a modified ChIP-seq study in *Pseudomonas aeruginosa* reports approximately 180 nascent transcripts that contain mostly non-overlapping enrichment peaks for both RsmA (a CsrA homolog) and Hfq (out of 560 total RsmA enrichment peaks) (**Gebhardt et al., 2020**). These nascent transcripts are likely simultaneously bound by both regulators, as most of these peaks do not directly overlap. Given the ternary CsrA-sRNA-complexes we observed *in vitro* (Figure 2) and the high throughput studies mentioned above (**Caillet et al., 2019; Gebhardt et al., 2020**), the possibility of sRNA-mediated CsrA-sRNA-Hfq ternary complexes forming in our *in vivo* assays, and, more broadly, endogenously throughout *E. coli* post-transcriptional regulation seems likely. The bulky ternary complexes could have inhibited *in vivo* detection of some of our CsrA-sRNA pairs in the context of our GFP-complementation assays (Supplementary Figure S4).

Overall, from a Csr systems perspective, remodeled sRNA-mRNA regulation allows inference of a new paradigm for understanding the scope of Csr network regulation. Prior omics studies of the Csr system imply wide-spread effects of CsrA deletion, more than what can be accounted for by the number of confirmed mRNA targets (**Potts et al., 2017; Sowa et al., 2017**). While CsrA regulation of transcription factors and sigma factors, such as *nhaR, sdiA*, *rpoE*, and *iraD* (an anti-adaptor protein that inhibits RpoS degradation) (**A Pannuri et al., 2012; Park et al., 2017; H. Yakhnin et al., 2017; H. Yakhnin, Baker, et al., 2011**), begin to explain these effects, crosstalk between CsrA and post-transcriptional sRNA regulators is likely to contribute. However, these crosstalk effects are difficult to extract from high throughput RNAseq and proteomics studies because the effects of CsrA remodeling sRNA regulatory networks are likely, as we propose, target specific. We anticipate that an *in vivo* role for CsrA in remodeling sRNA-mRNA regulation extends beyond the examples highlighted here. While precise mechanistic characterization is still needed, CsrA-remodeled sRNA-mRNA regulatory interactions may serve as templates for better understanding how global RNA-binding proteins extend their regulatory reach in cells. Even more broadly, this work sets the stage to investigate the full physiological impacts of the overlapping network interactions between CsrA and other global RBP networks.

## Materials and Methods

### Bacterial strains and growth conditions

All *E. coli* strains used in experiments are derivatives of K-12 MG1655 and are described in Supplementary Table S1S4. DH5α (NEB) and BL21 (DE3) (NEB) *E. coli* strains were used for cloning and protein purification, respectively. Routine cultures were performed in LB medium (Miller) (BD Biosciences) with 50 µg/mL kanamycin and/or 100 µg/mL carbenicillin as required. Starter cultures were grown from single colonies overnight in 5 mL LB (test tubes) at 37°C with 200 rpm orbital shaking. Cultures were diluted 1:100 in LB media for fluorescence assays and protein purification except where noted otherwise. Final concentrations of 1 mM isopropyl β-D-1-thiogalactopyranoside (IPTG) and 100 ng/mL anhydrotetracycline (ATc) were used in cultures to induce plasmid expression. Congo red plate assays were performed on agar plates containing LB medium without salt (10 g/L yeast extract, 5 g/L tryptone, 0 g/L NaCl), 40 µg/mL Congo red dye, 20 µg/mL Coomassie Brillant Blue G 250 dye, 50 µg/mL kanamycin, and 1 mM IPTG (Bak et al., 2015). Kanamycin resistance cassettes were cured from *sRNA::kan* deletion strains (Supplementary Table S1S4) and confirmed by colony PCR prior to use as previously described (Cherepanov & Wackernagel, 1995; Datsenko & Wanner, 2000).

### Plasmid construction

Plasmids used in this study are documented in Supplementary Table S1S4. All plasmids were constructed by Gibson assembly or purchased from Genscript. Oligonucleotide primers for Gibson assembly were designed using the NEBuilder 2.0 web tool and are included in Supplementary Table S2S6. DNA oligonucleotides and double stranded DNA fragments (GBlocks) were purchased from Integrated DNA Technologies. Plasmids were verified by Sanger sequencing (University of Texas GSAF core). Details on cloning methods used for construction of pTriFC plasmids, pHL600 pLacO-*sRNA* pTetO-*csrA* plasmids, pHL1756 5’UTR-*gfp* reporter plasmids, pBTRK-pLacO-sRNA plasmids, and pBTRK-pLacO-*fucP-gfp* (CmR) plasmids are included in Supplementary Methods. It should be noted that the pBTRK-pTrc-Empty plasmid, a generous gift from the lab of Dr. Brian Pfleger that contains a very low copy number pBBR-1 origin (∼1-3 copies per cell) (Hernández Lozada et al., 2018; Youngquist et al., 2013), was used as a parent plasmid for the pBTRK-pLacO plasmids constructed in this work.

### Strain construction

A previously-demonstrated CRISPR-cas9 genome modification protocol was used to construct *Δspf* and GGA:GCA mutant *spf* K-12 MG1655 strains (Mehrer et al., 2018). It should be noted that the *spf* gene encodes the Spot 42 sRNA. This approach uses a two-plasmid system to achieve genomic deletion or insertion. The first plasmid, a generous gift from the lab of Dr. Brian Pfleger, provides kanamycin-resistant guide RNA (gRNA) expression under the constitutive J23119 promoter and is termed pgRNA. The second plasmid, pMP11, is a temperature sensitive, carbenicillin-resistant plasmid that contains (1) wild type cas9 from *Streptococcus pyogenes* under the J23119 constitutive promoter, (2) λ-red recombinase under the pBAD promoter, and (3) a gRNA specific to the pgRNA plasmid to enable its removal under a pTeto promoter.

A gRNA for deletion of the *spf* gene was designed using the CRISPR gRNA Design webtool from Atum (atum.bio). This sequence was cloned into the gRNA plasmid by Gibson Assembly (Supplementary Table S2S6). Additionally, a dsDNA gBlock was designed to contain the desired *spf* deletion (here, the last 7 nt of the sRNA) or mutation sequence plus 500 base pairs of sequence homologous to the genome upstream and downstream of the edit site. To avoid gRNA-directed cleavage of the dsDNA homology GBlock, we targeted the gRNA to the CDS of a protein-coding gene, *yihA*, 150 nucleotides 3’ of *spf*. Synonymous codon mutations were made to the gRNA binding site in the dsDNA homology GBlock to protect it from pgRNA-directed cleavage. As a result, the 500 nucleotide homology regions were defined to be 500 nucleotides 5’ of *spf* and 3’ of *yihA* (Sequence in Supplementary Table S2S6).

K-12 MG1655 *E. coli* containing the pMP11 plasmid was grown in 5 mL of LB overnight to saturation. The overnight culture was diluted 1:100 in 50 mL of SOB media (BD Biosciences) and grown at 30 °C until an OD_600_ of 0.2. The λ-red genes were then induced by addition of 1% w/v L-arabinose. At an OD_600_ of 0.4-0.6, the culture was made electrocompetent according to common lab procedures. 50 ng of the gRNA plasmid and 25-50 ng of the dsDNA homology block was added to a single aliquot of electrocompetent cells, and subsequently transformed. Transformants were recovered in 1 mL of SOB media at 30 °C for at least three hours, plated onto LB agar plates containing kanamycin and carbenicillin, and grown at 30 °C. Gene deletions were confirmed via colony PCR (Supplementary Table S2S6) and sanger sequencing of the purified pcr fragment.

Upon verification of the *spf* deletion or mutation, the gRNA plasmid was cured by growing a single colony in 5 mL of LB containing carbenicillin and 0.2 ng/mL aTc overnight at 30 °C. Single colonies were of this culture were generated and streaked on carbenicillin versus kanamycin and carbenicillin LB agar plates to confirm pgRNA plasmid removal. The pMP11 plasmid was then cured by patching successful colonies onto LB agar only plates and growing overnight at 42 °C. Removal of pMP11 was verified by streaking on LB agar only versus LB agar plates with carbenicillin.

### Analysis of Csr RNAseq data

RNA sequencing and downstream data analysis was previously performed for *csrA::kan* mutant, Δ*csrB* Δ*csrC*, and wild type (WT) K-12 MG1655 *E. coli* strains before and after glucose deprivation in M9 minimal media (Sowa et al., 2017). Briefly, biological triplicate cultures were grown to an OD_600_ of 0.6 in M9 media with 0.2% glucose before centrifugation and resuspension in fresh M9 media lacking glucose. Samples were taken 10 min and 0 min prior and 30 min, 60 min, 180 min, and 300 min after glucose starvation. Aligned, counted reads from the previous work were reanalyzed by DESeq2 to include sRNAs and tRNAs (rRNA read counts were still excluded from analysis). As done in the prior work, all samples of the Δ*csrB* Δ*csrC* strain collected 300 min post stress were excluded from further analysis due to large variation amongst the replicas. Additionally, principal component analysis identified one sample, a replica of the of Δ*csrB* Δ*csrC* strain collected at time 0, that clustered far from the rest of the data. Data from this sample were also excluded from further analysis. Strain and time factors were paired in order to identify differentially expressed genes between the WT and *csrA::kan* strains and the WT and Δ*csrB* Δ*csrC* strains at each time point. sRNAs differentially expressed (log2 fold change > 1 or < -1 and Wald test P-adjusted < 0.05) between the WT and Δ*csrB* Δ*csrC* strains at two time points or more were considered as potentially interacting with the Csr system and selected for further study (Supplementary Table S3S7). It should be noted that by these criteria, no sRNAs were differentially expressed between the WT and *csrA::kan* strains. As levels of free CsrA are likely increased in the Δ*csrB* Δ*csrC* strain (Sowa et al., 2017), we considered differential expression between the WT and Δ*csrB* Δ*csrC* strains to indicate sRNAs likely impacted by CsrA.

### Thermodynamic prediction of sRNA binding sites

The biophysical model estimates the free energy of CsrA binding to pairs of 5 nucleotide potential binding sites within an RNA of interest based on RNA sequence, structure, and inter-site spacing. Based on these predictions, we analyzed the 15 most likely, i.e., lowest free energy, CsrA-RNA binding conformations, to identify putative CsrA-RNA binding regions. Previously, we confirmed the utility of this model for correctly predicting CsrA binding sites for 6 mRNAs of 8 total that have documented CsrA footprints. Within this subset, the model also successfully captured CsrA footprints for 3 mRNAs that lack the preferred GGA motif. When the model was run for sRNAs, predicted CsrA binding sites overlap GGA motifs found within the sRNA sequences (Supplementary Table S4S3). They also highlight non-GGA containing regions that may contribute, albeit likely transiently, to sRNA-CsrA recognition and binding in these sRNAs. Importantly, predicted sRNA-CsrA binding sites in the FnrS and MicL sRNAs, which lack GGA motifs, highlight a distinct region in each sRNA that contains a bulk of the predicted binding sites. CsrA-sRNA binding sites were predicted as previously done for mRNAs (Leistra, Gelderman, et al., 2018) and are further explained in the Supplementary Methods.

### Protein purification

CsrA was purified as previously published (Dubey et al., 2005) with minor modifications, as described in the Supplementary Methods. Hfq was purified according to published methods (Santiago-Frangos et al., 2016) and generously gifted to us by the lab of Dr. Sarah Woodson.

### *In vitro* electrophoretic mobility shift assays

Primers were designed to amplify sRNAs (Supplementary Table S2S6) with an upstream T7 promoter from genomic K-12 MG1655 DNA (wild type sRNA sequences) or previously constructed plasmids (mutant sRNA sequences). Forward primers were designed to contain four random nucleotides (GACT) upstream of the promoter sequence (TAATACGACTCACTATAGGGAGA). PCR product sizes were verified on a 1% agarose gel prior to PCR clean up (DNA Clean & Concentrator-5, Zymo). A DNA fragment for transcription of *lpp* (*lpp* sequence preceded by aforementioned T7 transcription-enabling sequence) was manufactured by Integrated DNA Technologies (Supplementary Table S2S6).

*Lpp* was *in vitro* transcribed using the MEGA Script IVT Kit (Thermo Fisher Scientific) according to manufacturer instructions (with incubation time extended to 6 hr as recommended for < 500bp RNA products). sRNAs were transcribed with modifications to enable internal P-32 labeling. Specifically, 1.5 µL UTP [α-32P] (3000 Ci/mmol 10 mCi/ml, 500 µCi, PerkinElmer) was used to replace equivalent unlabeled UTP from the MEGA Script IVT kit. Following MEGA Script T7 DNase digestion, RNA recovery was performed using RNA Clean & Concentrator-5 (Zymo Research) according to manufacturer instructions. RNA samples were re-suspended in 20 µL of nuclease-free water following recovery and, for sRNAs only, free NTPs removed using DTR Gel Filtration Cartridges (EdgeBio) following manufacturer instructions. RNA concentration was measured via spectrophotometry, and, in the case of *lpp,* transcript quality validated on an 8% urea gel, run at 100V for 3 hours, and stained with Sybr Green II (Thermo Fisher Scientific).

EMSAs were performed largely according to the TBE CsrA-RNA gel shift protocol detailed in (A. V Yakhnin et al., 2012), with modifications to support the use of a different CsrA dilution buffer (Dubey et al., 2005). Twelve µL binding reactions were comprised of 10 nM denatured sRNA (3 min at 85 °C), 1 µL 10X CsrA binding buffer (100 mM Tris-HCl pH 7.5, 100 mM MgCl_2_, 1 M KCl), 4 µL CsrA dilution buffer (same as CsrA storage buffer: 10mM Tris-HCl, 100 mM KCl, 10 mM MgCl_2_, and 25% glycerol, pH 7.0), 11% glycerol, 20 mM DTT, 333 U/mL RNase Inhibitor, Murine (NEB), 1.5 µg/µL yeast total RNA, 0-6 μM CsrA, and 0, 0.375, 1.5, or 3 μM Hfq. These concentrations of Hfq were selected to mimic the expected relative *in vivo* expression ratios of CsrA to Hfq based on estimations of their *in vivo* concentrations: 0.4-10 µM hexameric Hfq per cell (Kajitani et al., 1994; Moon & Gottesman, 2011; Wagner, 2013) and 6-17µM dimeric CsrA (Romeo et al., 2013). Importantly, each reaction contained an excess of total yeast RNA (Thermo Scientific) to exclude results due to non-specific binding. For these experiments, radiolabeled sRNAs (10 nM) were incubated with 3 µM or 6 µM concentrations of CsrA at 37 °C for 30 min prior to loading and running on a 10% non-denaturing polyacrylamide gel (A. V Yakhnin et al., 2012) with 0.5X TBE running buffer (IBI Scientific, 10x composition: 89 mM Tris, 89 mM Boric Acid, 2 mM EDTA) at 170 V between 5 hours and overnight (depending on sRNA) at 4 °C. These concentrations of CsrA represent Protein:sRNA ratios of 300:1 and 600:1, respectively. These ratios were selected to screen for CsrA-sRNA binding based on CsrA binding affinities reported for Spot 42, GadY, GcvB, and MicL, interactions presumed to be relevant *in vivo* based on significant enrichment in CsrA CLIP-seq data (Potts et al., 2017). Briefly, these previous *in vitro* binding assays employed molar ratios of CsrA:sRNA ranging from 50:1 to 2,000:1 (5 nM CsrA:0.1 nM sRNA to 200 nM CsrA:0.1 nM sRNA) and determined dissociation constants in the range of 100:1 to 660:1 CsrA:sRNA (10 nM CsrA:0.1 nM sRNA to 66 nM CsrA:0.1 nM sRNA). While our assays scale up the total molar amount of each component, they test CsrA-sRNA binding at molar ratios of 300:1 and 600:1 CsrA:sRNA. These ratios are within the range of CsrA-sRNA binding affinity presumed to be physiologically relevant in (Potts et al., 2017). Additionally, all binding assays are performed in large excess of yeast total RNA (88ng/µL, as in (A. V Yakhnin et al., 2012)) to inhibit non-specific association of CsrA to labeled sRNAs.

Gel exposure on a bioWORLD bioExposure cassette was phosphor-imaged at 1000V using Typhoon FLA 700 (GE Health Life Science).

MicL-*lpp*-CsrA EMSA was performed with slight modifications. Larger reaction volumes (15 uL) were used to accommodate addition of the *lpp* 5’ UTR (80 or 240 nM final concentration) and relative molar amounts of sRNA and CsrA equivalent with the gels described above were included (8 nM denatured sRNA and 0, 2.4, or 4.8 μM CsrA). All other components of the binding reaction were scaled appropriately.

### Tri-Fluorescence Complementation assays

Fluorescence complementation assays to detect RNA-protein binding *in vivo* were conducted as previously described, with modifications as described in the Supplementary Methods.

### GFP fluorescence quantification

#### sRNA-CsrA sponge activity

Plasmid-based screening of sRNA-CsrA sponge activity was inspired by a previous study (Adamson & Lim, 2013; Leistra et al., 2017). Here the plasmid encoding inducible CsrA and sRNA expression was altered to control expression of the sRNA with the stronger pLacO promoter (rather than a pTetO promoter used in the previous study). Expression of CsrA was controlled with the weaker pTetO promoter (rather than the pLacO promoter used in the previous study). The modified plasmid system was expressed in a K-12 MG1655 *E. coli* strain deleted for the Csr system (Δ*csrA* Δ*csrB* Δ*csrC* Δ*csrD*) and lacking the *pgaABCD* and *glgCAP* operons, as before (Adamson & Lim, 2013; Leistra et al., 2017). Single colonies of Δ*csrA* Δ*csrB* Δ*csrC* Δ*csrD* Δ*pgaABDC* Δ*glgCAP lacI_q_* K-12 MG1655 *E. coli* cells (Adamson & Lim, 2013) harboring pHL600 pLacO-sRNA pTetO-*csrA* and pHL1756 *glgC-gfp* plasmids were grown up overnight in 5 mL cultures. Biological triplicate overnights were seeded in 30 mL of LB (1:100 dilution, in flasks) with 100 ng/mL aTc to induce CsrA expression. After growth to an OD_600_ of 0.4 (∼ 3 hr), each culture was split in half and 1 mM IPTG was added to one 15 mL volume to induce sRNA expression. Green fluorescence was measured with a BD FACSCalibur flow cytometer (∼25,000 cells per sample) after 2 additional hours of growth to an OD_600_ of 1.5-2.0. Fold change in median green fluorescence between +sRNA and –sRNA conditions was determined for each biological replicate and averaged (n = 3). Two-tailed heteroscedastic t-tests were used to assess whether fold change in *glgC*-*gfp* fluorescence upon sRNA induction was significantly increased relative to that of the random RNA (P-value < 0.05). The assay was similarly conducted in the absence of induced CsrA expression, with the exception that each biological replicate starter culture was seeded in 15 mL of LB in technical duplicate. Once cultures reached an OD_600_ of 0.4, sRNA expression was induced in half of the cultures with 1 mM IPTG such that a *+*sRNA and –sRNA culture was obtained for each biological replicate. Green fluorescence was measured at the same conditions. It should be noted that in the absence of CsrA induction, cultures grew more slowly: 4.5 hr to OD_600_ 0.4 and approximately 4 additional hours to OD_600_ 1.5.

#### Impact of CsrA on sRNA-mRNA regulation

The same *E. coli* strain and plasmid system were used to assay the impact of CsrA on sRNA-mRNA regulatory pairs. Here, the 5’ UTRs of mRNAs regulated by sRNAs were used as translational GFP fusion reporters, as is typically done to test sRNA-mRNA interactions *in vivo* (Beisel & Storz, 2011; Bobrovskyy et al., 2019; Miyakoshi et al., 2019). Briefly, 5’ UTRs were defined as the region between the transcription start site and the mRNA start codon. This region, plus the first 100 nucleotides of mRNA coding sequence, was cloned in frame with *gfp* as previously published (Leistra, Gelderman, et al., 2018; Sowa et al., 2017). For simplicity, the 5’ UTR term is used to refer to the whole region cloned in frame with *gfp*. For mRNAs that code for membrane proteins, only the first 10 nucleotides of coding sequence were included in the reporter to minimize the impact highly hydrophobic amino acids might have on proper GFP folding and function (see Supplementary Table S1 S4 for full sequences tested). This length of sequence was chosen to reflect the range of coding sequence that is occluded upon ribosome-RBS binding (Espah Borujeni et al., 2017). However, if the sRNA binding site is or overlaps 3’ of the first 10 coding sequence nucleotides, the 5’ UTR was extended to include 10 nucleotides of coding sequence nucleotides beyond the 3’ edge of the sRNA binding region. For mRNAs in which the transcription start site is not known, the 100 nucleotides preceding the start codon was designated as the 5’ UTR (Sowa et al., 2017).

Single *E. coli* colonies expressing a pHL600 pLacO-sRNA pTetO-*csrA* plasmid and a corresponding pHL1756 5’ UTR*-gfp* plasmid were grown in 5 mL LB overnight to saturation. Saturated starter cultures were diluted 1:100 into 15 mL of LB (in flasks) four times. After growth to an OD_600_ of 0.2-0.3, sRNA and CsrA expression was induced (1 mM IPTG and 100 ng/mL ATc, respectively) such that the following four conditions were obtained for each biological replicate: – sRNA –CsrA; +sRNA –CsrA; –sRNA +CsrA; +sRNA +CsrA. Green fluorescence was measured with a BD FACSCalibur flow cytometer at several time points after induction. Fold change in median fluorescence value was calculated between induction conditions and averaged for biological replicates (n = 3). Paired two-tailed t-tests (P-value < 0.05) determined whether changes in fluorescence were significant. Heteroscedastic two-tailed t-tests (P-value < 0.05) were used to compare fluorescence fold changes measured for wild-type and mutant versions of the Spot 42 sRNA and *fucP*-*gfp* 5’ UTR reporter.

#### RNA extraction

Total RNA was extracted from cultures tested in GFP fluorescence assays by the TRIzol manufacturer protocol (Thermo Fisher Scientific) with a few modifications as described in the Supplementary Methods.

#### Northern blot analysis

Northern blotting was performed as previously reported (Cho et al., 2014), with a few modifications as described in the Supplementary Methods.

#### Enzymatic probing of CsrA interaction with Spot42 sRNA

RNase T1 probing of Spot42 was performed as previously described (Salvail et al., 2010) with a few modifications described in the Supplementary Methods.

#### Western blot analysis

To quantify CsrA expression levels in the GFP fluorescence assays, 9-10 mL of the 15 mL culture tested was pelleted by 10 min of centrifugation at 4000 rpm and 4°C. The supernatant was discarded and the pellet was flash frozen using liquid nitrogen and stored at -80°C for future use. Pellets were lysed and western blots conducted as described in the Supplementary Methods.

#### In vitro transcription and translation assays

Coupled transcription-translation assays were carried out with the PURExpress kit (New England BioLabs) as described in (**Lukasiewicz and Contreras, 2020**) with a few modifications described in the Supplementary Methods.

#### qRT-PCR quantification of sRNA effects on CsrA mRNA targets

Potential sRNA sponge or sequestration activity for endogenous CsrA and its mRNA target network was tested in K-12 MG1655 strains. CsrB, McaS, or SgrS sRNAs were cloned into the low copy number pBTRK-pLacO-Empty parent plasmid and expressed in a corresponding sRNA deletion strain (*ΔcsrBΔcsrC*, *ΔmcaS*, or *ΔsgrS* K-12 MG1655, respectively). Single colonies of these strains were grown up overnight in 5 mL cultures. Biological triplicate overnights were seeded in 30 mL of LB (1:100 dilution, in flasks). After growth to an OD_600_ of 0.2, each culture was split in half and 1 mM IPTG (final concentration) was added to one 15 mL volume to induce sRNA expression. Thirty minutes later, at an OD_600_ of 0.6, 5 mL volumes of all samples were harvested by centrifugation. RNA extraction was performed as described above. After verifying RNA quality, 10 µg of RNA was treated with DNase (DNAse I RNas-free, NEB) for 10-15 minutes at 37 °C. RNA was immediately re-purified with spin columns (RNA Clean and Concentrator-5 kit, Zymo Research) and concentration determined by nanodrop.

Abundance of the following RNAs was assessed by qRT-PCT in each sample: CsrA target mRNAs *glgC* and *pgaA;* non-target mRNA *phoB;* the sRNA induced, i.e., CsrB, SgrS, or McaS; and housekeeping reference RNAs *secA* and 16s. The *secA* mRNA and the 16s rRNA were both employed as housekeeping references as the endogenous mRNAs in question were much lower in abundance than the standard 16s rRNA reference. Additionally, *secA* has been previously used a housekeeping reference target in prior Csr study (Butz et al., 2019). Primers for qRT-PCR reactions were designed with the IDT PrimerQuest tool, specifying dye-based qPCR quantification. All primer pairs yielded amplicons of 75-125 nucleotides. Primer efficiencies were determined for each pair to be 90-105% across a 10^4^-fold range of RNA concentrations (0.005 to 5.0 ng or 0.05 to 50 ng RNA). It should be noted that 50 ng of RNA was needed to detect mRNAs with an approximate threshold cycle (C_T_) value of 20, while only 0.005 ng was needed to detect sRNAs and the 16s rRNA at the same approximate C_T_ value.

Biological triplicate RNA samples were tested in technical triplicate for each RNA target (i.e., each primer pair). All qRT-PCR reactions were performed with the Luna Universal One-Step RT-qPCR kit (E3005 NEB) according to manufacturer protocol with a few modifications. Reactions were prepared as follows in 384 well plates (MicroAmp Optical 184-Well Reaction Plate, Thermofisher): 5 µL Luna one-step reaction mix, 0.5 µL Luna RT mix, 0.4 µL of 10 µM forward primer, 0.4 µL of 10 µM reverse primer, 1µL of 0.005 ng/µL or 50 ng/µL RNA sample, and RNase free water up to 10 µL. qRT-PCR reactions were performed with ViiA7 thermocyclers (Applied Biosystems) according to Luna Universal One-Step RT-qPCR kit recommendations. Default melt curve settings for the thermocycler were used.

Control reactions lacking RNA were performed for each primer pair and reactions lacking the RT mix were performed for each RNA sample. After ensuring detection of *glgC*, *pgaA*, *phoB*, and *secA* mRNA targets in 50 ng RNA and CsrB, McaS, SgrS and 16s RNA targets in 0.005 ng RNA with single-peak melt curves, the ΔΔC_T_ method was used to determine relative change in RNA level upon sRNA induction. ΔC_T_ values were calculated for mRNAs relative to the *secA* mRNA reference and for sRNAs relative to the 16s rRNA reference for each biological replicate. Paired two-tailed t-tests between uninduced and induced ΔC_T_ values of a given RNA target were used to determine significant changes in RNA abundance (upon induction of a sRNA). ΔΔC_T_ values were calculated, averaged across biological replicas, and presented as log2 fold changes.

#### Congo red plate assay

Wild type and cured Δ*mcaS* K-12 MG1655 *E. coli* strains were made competent according to lab protocols. Wild type and mutant McaS (5’-most GGA:CCA mutant, Supplementary Table S1S4), cloned in the low copy number pBTRK-pLacO-Empty plasmid (Youngquist et al., 2013), were transformed into the Δ*mcaS* strain. An empty control pBTRK-pLacO-Empty plasmid was transformed into both the wild type and Δ*mcaS* strains. Single colonies were grown up in biological triplicate 5 mL LB cultures. 10 µL of each saturated overnight culture was spread on Congo red agar plates and grown at room temperature for 36-48 hours. Plates were placed on a UV to white light conversion screen (BioRad) and backlit with LED lighting for imaging.

#### Fluorescence assays with induction of genomic Spot42

To evaluate the biological relevance of CsrA impacting the Spot 42-*fucP* interaction, culturing conditions were developed to allow native expression of the Spot 42 sRNA in K-12 MG1655 *E. coli* to regulate *fucP*-*gfp* reporters expressed from the low copy number pBTRK-pLacO (CmR) plasmid. The media utilized was M9 Media supplemented with 0.4% glycerol, 0.2% casamino acids, 10 µg/mL thiamine, 2 mM MgSO4, and 0.1 mM CaCl2, which has been previously demonstrated to allow for high genomic expression of the Spot 42 sRNA upon bolus addition of 0.2% glucose (Beisel & Storz, 2011). For the fluorescence assay, the pBTRK-pLaco-*fucP-gfp* (CmR) plasmids containing wild type and mutant *fucP* 5’ UTRs with disrupted Spot 42 binding sites (Supplementary Table S1S4) were transformed into wild type and *Δspf* K-12 MG1655 *E. coli* strains. Single colonies were grown overnight in 5 mL of the M9 Media in biological triplicate until saturation and seeded for culturing by diluting 1:100 into 25 mL of the M9 Media. Additionally, the *fucP*-GFP reporter was induced at seeding using IPTG at a final concentration of 1 mM. Bolus glucose was added to a final concentration of 0.2% at an OD600 of 0.2, approximately 3.5 hours after seeding. The cultures were then sampled at various timepoints following induction and fluorescence was measured using a BioTek Cytation3 with an excitation wavelength of 488 nm and an emission wavelength of 513 nm. Absorbance at 600 nm was also measured and the fluorescent values were normalized to control for differences in number of cells per well. Average fluorescence values of each *fucP-gfp* reporter were compared between wild type and *Δspf* strains with heteroscedastic two-tailed t-tests (n = 3, P-val < 0.05) to determine significant regulation by genomic Spot42. To compare Spot42 regulation of different *fucP-gfp* reporter constructs, fold change in fluorescence for each construct was determined as the ratio between average median fluorescence in the wild type and *Δspf* strains (n = 3). Heteroscedastic two-tailed t-tests were utilized to determine if fluorescence fold changes were significantly different (P-val < 0.05) between the *fucP-gfp* constructs.

## Data Availability

Predicted CsrA-sRNA binding sites data are included in the SI. Re-analyzed transcriptomics data were previously published in (Sowa et al., 2017).

## Acknowledgements

We thank Dr. Sarah Woodson, Dr. Ewelina Małecka, and Dr. Penny Peng for helpful conversations and a generous gift of purified Hfq protein. We also thank Dr. Susan Gottesman, Dr. Tony Romeo, and Dr. Han Lim for their gifts of several *E. coli* strains. Acknowledgment is also given to Dr. Brian Pfleger for sharing plasmids (pBTRK-pTrc-Empty, pMP11, and pgRNA), Dr. Svetlana Harbaugh for providing us with plasmids for the IVTT assays (pUC19-sfGFP) and to Dr. Howard Salis for his assistance in running the previously published biophysical CsrA-RNA interaction model. We additionally thank Emily Bowman, Bridget Li, Matthew Law, and Rachael M. Cox, for their assistance in preliminary electrophoretic mobility shift assay work. We finally thank Jesus Garcia for his assistance in cloning and fluorescence data collection and other members of the Contreras group for helpful conversations.

