## Supplemental Figures and Methods for "CsrA Shows Selective Regulation of sRNA-mRNA Networks"

### Supplementary Methods

#### Plasmid construction

For pTriFC plasmids cloned in-house, backbone fragments were generated by restriction digest (AatII and SpeI) and inserts were PCR amplified from *E. coli* genomic DNA with homologous overlaps.

The pHL600 pLacO-*csrB* pTetO-*csrA* (pCML 2556) parent plasmid was cloned from the previously published pHL600 pTetO-*csrB* pLacO-*csrA* plasmid (pCML 877) (61) by two rounds of Gibson assembly. First, PCR amplification was used to generate a backbone fragment with homologous overlaps to a pLacO insert, which was generated by annealing complementary primers. After assembly and sequence verification, PCR amplification was used to generate a new backbone fragment with homologous overlaps to a GBlock sequence containing the pTetO promoter. After assembly and sequence verification, the resulting plasmid (pCML 2556) served as a parent for constructing the pHL600 pLacO-sRNA pTetO-*csrA* plasmids. The backbone and insert fragments were generated by PCR amplification of the parent plasmid and *E. coli* genomic DNA, respectively. sRNA sequences were defined as annotated in the NC000913.3 genome from NCBI (downloaded prior to fall 2018 annotation updates). Mutant sRNA sequences were purchased as GBlocks from IDT. 5' UTR-*gfp* translational fusion reporters were constructed by Gibson assembly in the pHL1756 parent plasmid (Supplementary Table S1 and S2) as previously described (55, 62).

The pBTRK-pTrc-Empty control plasmid, a generous gift from the lab of Dr. Brian Pfleger, contains a low copy number pBBR-1 origin (~1-3 copies per cell) (63, 64). This template was PCR amplified upstream of the pTrc promoter and downstream of the MCS. pBTRK-pLacO-Empty was assembled with this fragment and a GBlock containing the pLacO promoter sequence, MCS, and homologous overlaps. pBTRK-pLacO-Empty served as the parent for constructing the pBTRK-pLacO-sRNA plasmids. Restriction digest was used to generate the backbone fragment (SalI and SphI), while PCR amplification of the corresponding pHL600 pLacO-sRNA pTetO-*csrA* plasmid was used to generate sRNA insert sequences with homologous overlaps. Similarly, a modified version of the pBTRK-pLacO-Empty plasmid was used a parent plasmid for low copy number expression of the *fucP-gfp* reporter. The pBTRK-pLacO-Empty plasmid was modified for chloramphenicol resistance (rather than kanamycin) and to remove excess DNA 3' of the MCS and 5' of the rrnB terminator region by Gibson Assembly (Supplementary Table S2). This plasmid, pBTRK-pLacO (CmR), served as parent construct for inserting *fucP-gfp* translational reporter sequences. In most cases, the *fucP-gfp* sequence was PCR amplified from a corresponding pHL1756 5' UTR-*gfp* reporter plasmid with homologous overlaps to the pBTRK-pLacO-Empty (CmR) plasmid for Gibson Assembly. In other cases, a GBlock sequence of the desired insert sequence with homologous overlaps was purchased.

#### Design of mutant RNA sequences to confirm binding specificity

Mutations to RNA sequences were made to interrupt sRNA-CsrA interactions and, in the case of the Spot 42*,* sRNA-mRNA interactions. Mutant sequences tested are provided in Supplementary Table S1S4. In order to interrupt sRNA-CsrA binding, instances of the GGA nucleotide sequence, the known high-affinity CsrA-RNA binding motif, were taken as putative CsrA binding sites and mutated according to previous studies. GGA:CCA mutations were preferentially attempted in structural predictions (Vienna RNAfold webserver, default parameters) as this mutation has been shown to interrupt CsrA binding *in vitro* and *in vivo* (Patterson-Fortin et al., 2013; Vakulskas et al., 2016). A mutation was considered successful if (i) the predicted minimum free energy secondary structure remained the same as that of the wild type sRNA sequence, and if the (ii) free energy of folding as well as the (iii) predicted frequency of the MFE structure in the ensemble were consistent with those of the wild type sRNA sequence (± 1 kcal/mol was tolerated). If the GGA:CCA mutation was unsuccessful, GGA:GCA and GGA:CGA mutations were attempted, as these have also been shown to interrupt CsrA binding (Dubey et al., 2005). Compensatory mutations were made to nucleotides base-pairing with the GGA sequence in the wild type sRNA to maintain predicted structure if necessary. Here, GGA:UUA, GGA:GUA, or GGA:UGA mutations were made if a G nucleotide was predicted to base pair with a U nucleotide in the wild type structure (rather than a C). For FnrS, which lacks a GGA motif, CsrA binding sites predicted by the biophysical model were analyzed. Because stem loop 1 of the predicted wild type sRNA structure contains a bulk of the predicted CsrA binding sites (as well as the most likely pair of binding sites), two GG dinucleotides in the stem and two G nucleotides in the loop were mutated as follows: G:C and GG:CC, GC, or CG (termed "GGN Mutant", Supplementary Table S1S4). Here, an additional mutant FnrS sequence ("GG mutant 2," Supplementary Table S1) was made to eliminate an added, structured GGA sequence. It is also worth noting that two mutant McaS sequences were also constructed. The first interrupts both GGA motifs confirmed as CsrA binding sites *in vitro* ("dual GGA mutant," Supplementary Table S1S4), while the second mutant sequence interrupts only the 5'-most confirmed binding site ("5' GGA mutant," Supplementary Table S41).

Similar predicted secondary structure requirements were applied in design of Spot 42-*fucP* mRNA binding site mutations. Compensatory mutations previously made to Spot 42 and *fucP* 5' UTR in order to validate the Spot 42-*fucP* site 3 interaction via mutational reporter assays (Beisel et al., 2012) were determined to not impact predicted secondary structure of either RNA in our workflow and were tested *in vivo*. Similarly, mutation previously made to Spot 42 in order to validate interaction with other mRNA targets at site 1 was determined to not impact predicted Spot 42 secondary structure in our workflow and was tested *in vivo.* However, compensatory mutation to *fucP* at site 1 resulted in a minor alteration of predicted secondary structure. In order to maintain congruency with earlier literature, we constructed and tested this mutant in our mutational fluorescence reporter assays anyway. It should be noted, however, that we constructed an additional *fucP* site 1 mutant 5'UTR-*gfp* reporter in which predicted secondary structure was maintained (termed "site 1b mutant", see Supplementary Table S4). This construct similarly interrupted Spot 42-*fucP* activation in the absence of CsrA (Supplementary Figure S2).

**Protein purification**

CsrA-His6 was expressed from a pET-21a(+) vector (pCSB12) in BL21 DE3 E. coli. Saturated starter culture was diluted 1:100 into 500 mL of LB. The plasmid was induced with 1 mM IPTG after growth to an OD600 of 0.6 at 37°C. After 20 hours of additional growth at 37°C, the culture was split into two volumes and was harvested by centrifugation. Each pellet was washed with 200 mL 1X PBS and resuspended in 5 mL of lysis buffer (50 mM NaH2PO4, 300 mM NaCl, 10 mM Imidazole, 5 mM MgCl2 and 10% glycerol, adjusted to pH 8.0 with NaOH). Concentrated cell suspensions were combined and sonicated for 2.5 mins (10s on, 5s off, 10x) at a 40% power output setting (Q125 sonicator) 5 times. Soluble lysate was collected by centrifugation and purified by two rounds of nickel column purification.

Qiagen Ni-NTA agarose column resin was prepared with 5 mL of lysis buffer and drained according to manufacturer recommendations in a 5 mL polypropylene column (Qiagen). All soluble lysate (~10 mL) was loaded onto the column and rotated end over end for 1.5 hr at 4°C. The column was washed with 5 mL of wash buffer 1 (50 mM NaH2PO4, 300 mM NaCl, 5 mM

MgCl2, 20 mM Imidazole, 10% glycerol, adjust pH to 8.0 using NaOH) twice. Column-bound CsrA was eluted from the column in seven 1 mL volumes of elution buffer (50 mM NaH2PO4, 300 mM NaCl, 5 mM MgCl2, 250 mM Imidazole, 10% glycerol, adjust pH to 8.0 using NaOH). Purified fractions were analyzed by 15% SDS-PAGE and Coomassie staining (Supplementary Figure S1). All elution fractions were combined and exchanged into lysis buffer with 3kDa Ultra-4 centrifugal filter units (Milipore Sigma) at 4000 rpm and 4ºC. Protein was loaded onto a second Ni-NTA agarose column and the purification repeated with the following modification: the column was washed with 5mL of wash buffer 2 (50 mM NaH2PO4, 300 mM NaCl, 5 mM MgCl2, 50 mM Imidazole, 10% glycerol, adjust pH to 8.0 using NaOH) six times prior to elution. Purified fractions were analyzed by 15% SDS-PAGE and Coomassie (Supplementary Figure S1) Suitably pure fractions, i.e., those in which the intensity of the CsrA band was greater than 90% of the total lane intensity (assessed by imageJ), were combined and exchanged into CsrA storage buffer (10 mM Tris-HCl, 100 mM KCl, 10 mM MgCl2, 25% glycerol, pH 7.0). Protein concentration was determined by Bradford assay. Purified CsrA was diluted to 18 µM and stored at -20°C. It is worth noting that purified CsrA precipitated at higher concentrations in this storage buffer.

**Trifluorescence complementation assays**

Plasmids encoding CsrA-NYFP and MS2 binding domain (MS2BD)-sRNA fusions were constructed and expressed in Δ*csrB* Δ*csrC* Δ*pgaABDC* Δ*glgCAP* $lacI_q$ K-12 MG1655 *E. coli* cells (75) alongside a MS2 protein-CYFP fusion. As previously described, each sRNA of interest was tested with a negative control fusion lacking CsrA (i.e., empty-NYFP) (73, 74). Additionally, an MS2BD fusion of a specific random RNA sequence was tested with the CsrA-NYFP and Empty-NYFP fusions to control for indirect impacts of CsrA overexpression on cell growth. Biological triplicate colonies were picked and grown to saturation in 5 mL starter cultures at 37ºC overnight. Fresh 5 mL subcultures were seeded (1:100 dilution, in flasks) and grown at 37ºC until mid-exponential phase (OD 0.3-0.6). Expression of the fusion constructs from pLacO promoters was induced with 1 mM IPTG (final concentration). Cultures were then grown at 25ºC for an additional 22 hours. Yellow fluorescence of 100,000 single cells per sample was measured with a BD FACSCalibur and median fluorescence values were computed. For each MS2BD-sRNA fusion, fold change in average median yellow fluorescence values collected with the CsrA-NYFP fusion and the Empty-NYFP control fusion was determined. Fold changes were compared to that of the random RNA by two-tailed heteroscedastic t-tests. sRNAs with P-value < 0.05 were taken as significantly associating with CsrA *in vivo*. It should be noted that strains expressing plasmids with the SibA, SibD, and SibC sRNAs could not be consistently grown in either *E. coli* strain and were thus not tested in this assay.

**RNA extraction**

Total RNA was extracted from cultures tested in GFP fluorescence assays by the TRIzol manufacturer protocol (Thermo Fisher Scientific) with a few modifications. Briefly, 4-5 mL of culture was pelleted by centrifugation (4000 rpm, 4°C). The supernatant was discarded and the pellet was flash frozen with liquid nitrogen and stored at -80°C. Pellets were thawed on ice and resuspended in 1mL of room temperature TRIzol (Invitrogen) with 30s of rapid pipetting to homogenize the sample. After incubating at least 5 min at room temperature, 300 µL of chloroform/isoamyl alcohol 24:1 (Thermo Fischer Scientific) was added to each sample. Samples were inverted rapidly for 15s, incubated for an additional 3 min at room temperature, and centrifuged at 15,000 rpm for 15 minutes at 4°C. The clear upper aqueous phase was isolated without disturbing the red organic phase or cloudy interphase and transferred to RNAse-free 2.0

mL tubes containing 500 µL of ice-cold isopropyl alcohol, XX ammonium acetate, and 2.5 µL of GlycoBlue Co-precipitant (Thermo Fisher Scientific). RNA precipitated overnight (~16hours) at -20ºC and was pelleted by centrifugation at 4ºC (30 min 15,000 rpm). The supernatant was aspirated and pellets washed twice with 1 mL of 95% ethanol and once with 1 mL of 75% ethanol. Each wash was added without disturbing the pellet and briefly centrifuged (5 min 15,000 rpm at 4°C) prior to the next wash. Pellets were dried ~3min in an Eppendorf Vacufuge plus without centrifugation (aqueous vacuum setting only) or for ~30 minon a benchtop. RNA samples were rehydrated with 20-30 µL of RNAse free water and concentrations were determined with a Nanodrop spectrophotomer (Thermo Fisher Scientific). RNA quality was checked by running 1 µg of isolated RNA on a 1% agarose gel.

**Northern blot analysis**

Ladder (Promega ΦX174 DNA/HinfI dephosphorylated markers) and DNA oligonucleotide probes were 5' end-labelled for northern blotting with T4 polynucleotide kinase (NEB) as done previously (82). RNA samples (10 µg) were mixed with equal volumes of RNA loading dye (NEB), denatured for 5 minutes at 95ºC, and immediately loaded into a 10% polyacrylamide 7 M urea gel (SequaGel, National Diagnostics). RNA was separated by gel electrophoresis (BioRad Protean II) at 4ºC in 0.5X TBE buffer (IBI Scientific, 10x 89 mM Tris, 89 mM Boric Acid, 2 mM EDTA) for ~4.5hr at 12 watts. Separated RNA was then transferred to Hybond-N+ nylon membrane (GE Healthcare) at 4ºC for 18hr (15V) (BioRad Trans-Blot Cell) in 0.5X TBE buffer. After UV crosslinking (1200 µJ/cm2), membranes were hybridized in PerfectHyb Plus hybridization buffer (Sigma-Aldrich) with a radiolabeled DNA oligonucleotide probe at 42ºC for 3hr. An initial 20 min wash at 30ºC was performed in 5X SSC 0.1% SDS, followed by two 15 minutes washes in 1X SSC 0.1% SDS at 42ºC. After rinsing in Nanopure water, membranes were exposed to phosphor-screens for 12-48 hours and imaged with a Typhoon imager (GE Healthcare). Relative band intensities were determined using ImageJ.

**Western blot analysis**

Pellets were then thawed on ice and resuspended in 1 mL of 1x PBS. The samples were pelleted by centrifugation at 4000 rpm and 4°C. The supernatant was discarded and the pellets were resuspended in 1 mL fresh 1x PBS. Along with the frozen cell lysates from the fluorescence experiments, cultures of wild type K-12 MG1655 E. coli and ΔcsrA ΔcsrB ΔcsrC ΔcsrD ΔpgaABDC ΔglgCAP lacIq K-12 MG1655 E. coli (referred to as Δcsr below) were grown up to an OD600 of 1.5 and prepared following the same protocol.

Each sample was then sonicated on ice for 2.5 minutes: 10 seconds of sonication followed by 5 seconds of rest. This was done for a total of three rounds per sample. The samples were then centrifuged at 4000 rpm and 4°C. The supernatant, being the total soluble cell lysate, was aspirated and transferred to a clean 1.7 mL Eppendorf tube.

The total protein concentration of each cell lysate sample was measured using the Pierce Coomassie Plus (Bradford) Assay kit (Thermo Fisher Scientific). Protein gels were run using precast Mini-PROTEAN 16.5% polyacrylamide Tris-Tricine gel (Bio-Rad). For each sample, 5 µg of total soluble protein was diluted to a total volume of 10 µL with 1x PBS and mixed with an equal volume of 2x sample loading buffer (200 mM Tris-HCl pH 6.8, 2% SDS, 40% glycerol, 2% β-mercaptoethanol, ~ 0.5% w/v bromophenol blue). Samples were loaded after 5 min of denaturation at 95°C along with protein standards (Dual Xtra Prestained Protein Standard from Bio-Rad, Color Prestained Protein Standard Broad Range from NEB). Purified CsrA was also included to serve as a positive control for identifying the CsrA protein band in the subsequent western blot. The Tris-Tricine protein gel was run for 300 minutes at 70V (25°C) with 1X Tris-Tricine running buffer (100 mM Tris, 100 mM Tricine, 0.1% SDS). Separate protein samples were immediately transferred with the Trans-Blot SD Semi-Dry Transfer Cell (Bio-Rad) to a 0.2 µm Nitrocellulose Membrane (Bio-Rad) for 20min at 15V (25°C) with ice-cold transfer buffer (25 mM Tris, 192 mM Glycine, 20% v/v Methanol, 0.0075% w/v SDS) for western blotting.

Western blotting proceeded as previously described (54), with minor modifications. Following the transfer, the membrane blocked overnight in 5% milk TBS (w/v) solution at 4°C. The membrane was washed four times with 50 mL TBST (20 mM Tris, 500 mM NaCl, 0.05% v/v Tween20, pH 7.5) and once with 50 mL TBS (20 mM Tris, 500 mM NaCl, pH 7.5) for 5 minutes each (simply described as "washed" hereafter). The GroEL protein was blotted to serve as a loading control for one hour at room temperature with the anti-GroEL antibody (Novus Biologicals) at a 1:10,000 dilution in a 1% milk TBS solution. Following the incubation, the membrane was washed and the secondary incubation was done using an anti-Mouse HRP antibody (Promega) 1% milk TBS solution for one hour at room temperature. Following the incubation, the membrane was washed and visualized with the standard protocol of the Clarity Western ECL Substrate kit (Bio-Rad).

To continue blotting for further proteins, the membrane was washed and blocked for 3 hours at room temperature. To blot for CsrA, the membrane was washed and incubated overnight at 4°C using a 1:2500 anti-CsrA antibody (BiorByt) in 2.5% milk TBS (w/v) solution. After the primary incubation, the membrane was washed and the secondary incubation was done with a 1:1000 anti-rabbit HRP antibody (Abcam) in 1% milkTBS (w/v) solution for one hour at room temperature. Following an additional round of washes, the membrane was visualized again using the Clarity Western ECL Substrate kit (Bio-Rad). ImageJ was used to quantify band intensities for both CsrA and the loading control.

**Thermodynamic Predictions of sRNA Binding Sites**

This model estimates the free energy of a CsrA dimer binding a target RNA at a pair of potential binding sites based on target RNA nucleotide sequence and structure. Four free energy terms are estimated and summed to calculate a total free energy of CsrA-RNA binding: $\Delta G_{site\ 1}$, $\Delta G_{site\ 2}$, $\Delta G_{cooperativity}$, and $\Delta\Delta G_{unfolding}$. $\Delta G_{site\ 1}$ and $\Delta G_{site\ 2}$ are calculated with a position weight matrix (PWM) that estimates the free energy of CsrA binding to any 5 nucleotide RNA sequence based on similarity to the SELEX-derived AAGGA consensus sequence. The PWM was derived from experimentally-measured dissociation constants of five mutations to the AAGGA consensus sequence, one at each nucleotide position (Dubey et al., 2005). $\Delta G_{cooperativity}$ approximates the energetic contribution of inter-site spacing as a single CsrA dimer has been shown to simultaneously bind pairs site spaced 10-63 nucleotides (nt) apart *in vitro* (Mercante et al., 2009). Lastly, $\Delta\Delta G_{unfolding}$ quantifies the change in free energy of the mRNA unfolding required to accommodate CsrA binding at two 5 nt sites (Vienna RNA, Turner 1999 free energy parameters, 37°C, no dangling end energies) (Gruber et al., 2008; Lorenz et al., 2011). All possible pairs of CsrA-RNA binding sites with $\Delta G_{site} < 0$ are identified and the 15 most likely pairs, i.e., lowest total free energy of CsrA-RNA binding, are considered for analysis (see Supplementary Dataset S1 and Supplementary Table S4S3). Predicted secondary structures of non-CsrA-bound sRNAs were visualized with the *forna* RNA secondary structure visualization tool from Vienna RNA (Kerpedjiev et al., 2015). RNA sequence and predicted dot-bracket structure was supplied to the webtool. Additional minimum free energy prediction of the *fucP* mRNA 5' UTR used the Vienna RNAfold webserver with the same parameters (isolated base pairs were not avoided); results were similarly visualized with the *forna* webtool. A local installation of IntaRNA version 2.2.0 (Busch et al., 2008; Mann et al., 2017) was used to predict additional interaction sites between the Spot 42 sRNA and the 5'UTR of the *fucP* mRNA target. Default parameters were used, except for allowing multiple interactions to be predicted.

### Supplementary Figures

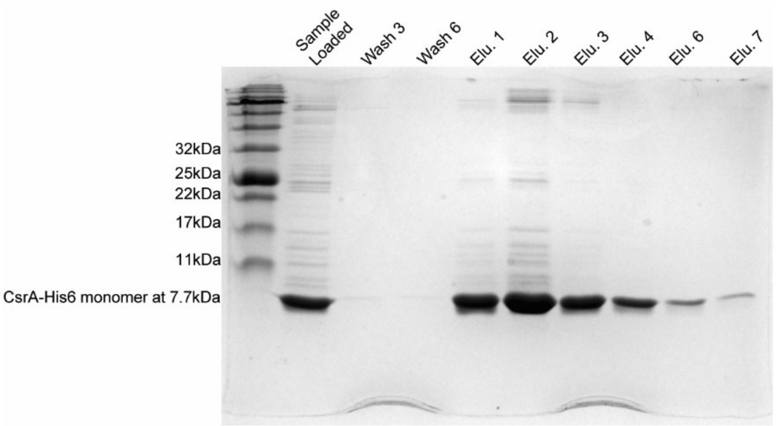


**Supplementary Figure S1. CsrA purification SDS-PAGE gel for *in vitro* EMSAs.** Stained with Coomassie. Named fractions from the second round of nickel column purification of CsrA-His6 were run on a 15% resolving gel (tris-glycine gel and buffer system). Elutions 1 and 3-7 were combined and transferred into storage buffer with 3kDa cut-off centrifugal filters.

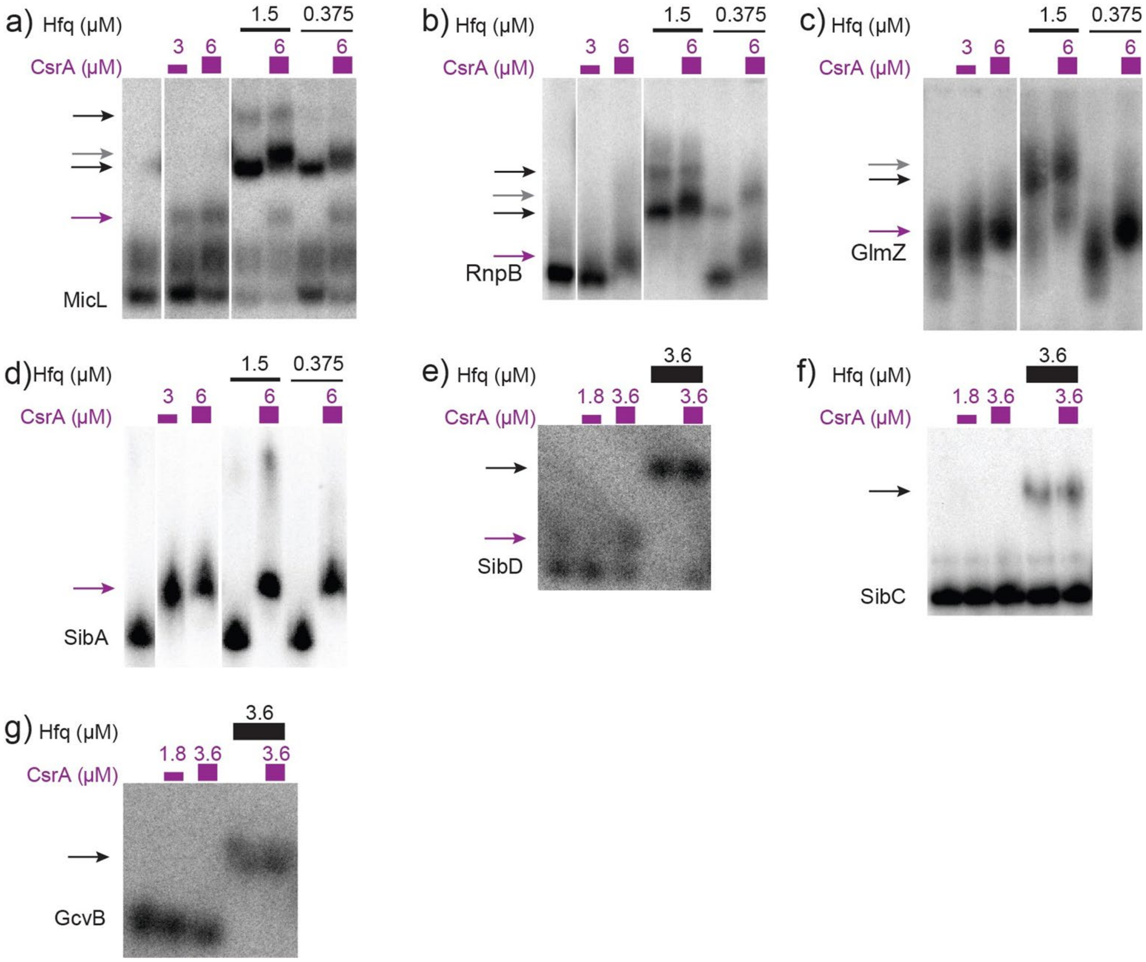


**Supplementary Figure S2. MicL, RnpB, GlmZ, SibA, and SibD bind CsrA *in vitro.*** (A-D) Radiolabeled sRNAs (10 nM) were incubated with increasing concentrations of purified CsrA (0, 3, or 6 μM) or Hfq (0, 0.375, or 1.5 μM) and separated on a 10% non-denaturing polyacrylamide gel. White space between lanes denotes non-contiguous lanes from the same gel. Complete EMSA gel images are included in Supplementary Figure S3. (E-G) Radiolabeled sRNAs (12 nM) were incubated with 1.8 and 3.6 μM CsrA or Hfq (0 or 3.6 μM) and separated as (A-D). (A-G) Purple arrows indicate shifts from sRNA-CsrA complexes (panels A –E). Black arrows indicate shifts detected from sRNA-Hfq complexes (A-C and E-G). Further shifts in sRNA-Hfq complexes upon inclusion of CsrA, i.e., likely CsrA-sRNA-Hfq ternary complexes, are shown as gray arrows (panels A-C).

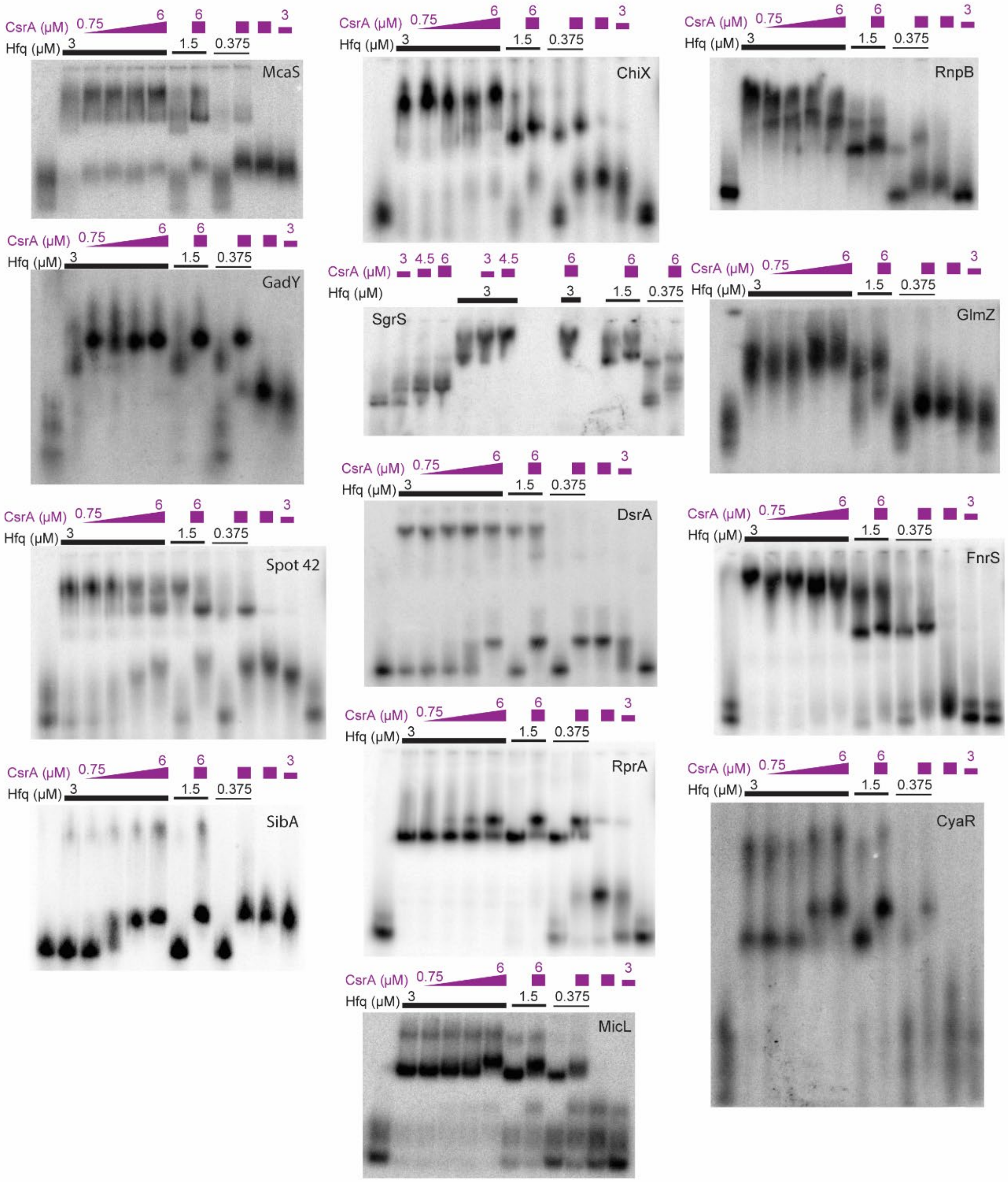


**Supplementary Figure S3. Complete images of EMSA gels** presented as non-contiguous lanes in Figure 2 and Supplementary Figure S2. Free RNA is the leftmost lane in each gel image (and, in some cases, is the rightmost lane as well). Protein concentrations are indicated above the lanes. The wedge of increasing CsrA concentrations is: 0.75, 1.5 uM, 3 uM, 6 uM across the four lanes covered by the wedge in each gel. It should be noted that the SgrS gel contains modified protein concentrations. These values are noted above each lane.

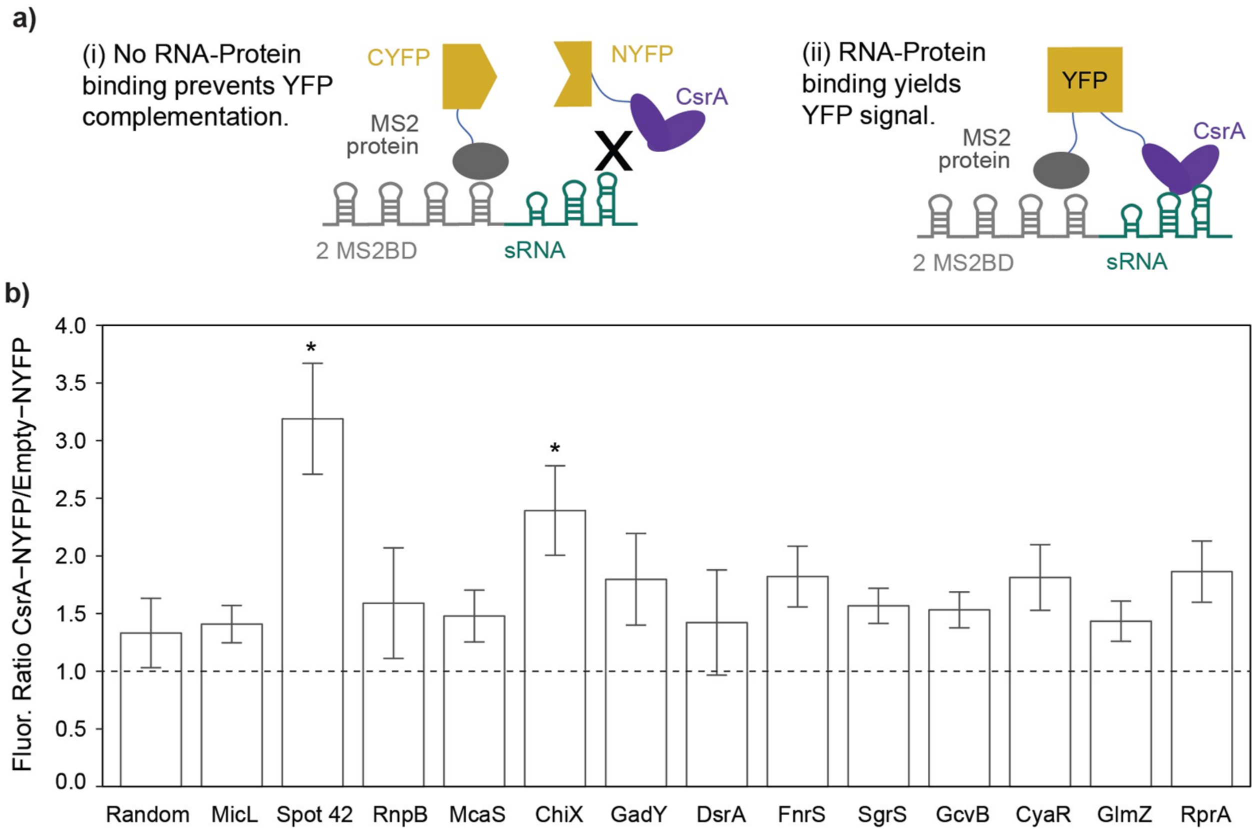


**Supplementary Figure S4. CsrA-sRNA binding detected in vivo by fluorescence complementation.** 4a) Three-part system for detecting sRNA-protein binding is comprised of: 2 repeats of the MS2 binding domain (MSBD) tagged 5' to an sRNA of interest, a MS2 binding protein-linker-CYFP protein fusion, and a CsrA-linker-NYFP protein fusion. Limited complementation in the absence of CsrA-sRNA binding (panel i) is differentiated from positive complementation in the presence of CsrA-sRNA binding (panel ii). 4b) Ratio of average yellow fluorescence measured for a sRNA with the full CsrA-NYFP fusion versus the Empty-NYFP control fusion in Δ*csrB* Δ*csrC* Δ*pgaABDC* Δ*glgCAP* $lacI_q$ K-12 MG1655 *E. coli*. Constructs were grown at 37ºC to approximately OD 0.5 before induction of all fusion constructs with 1 mM IPTG. Yellow fluorescence was measured after an additional 22 hr of growth at 25°C. Averages represent three biological replicates and error bars indicate propagated standard deviation. Asterisk indicates P-value < 0.05 of heteroscedastic T-test between CsrA-NYFP/empty-NYFP of an sRNA and that of the random negative control RNA.

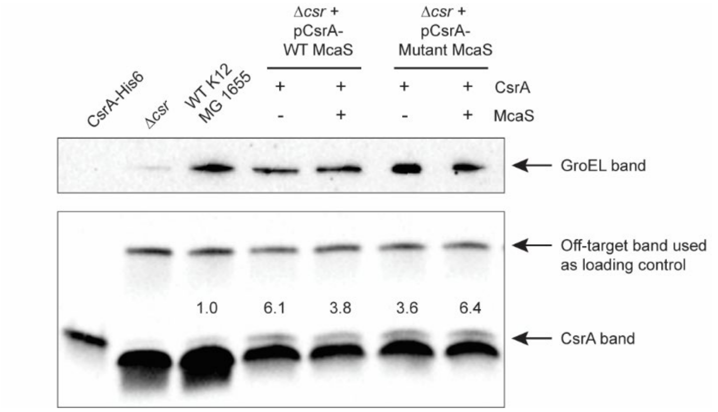


**Supplementary Figure S5. Western blot of overexpressed CsrA in g*lgC-gfp* reporter assays.** Named samples were run on a precast 16.5% Tris-tricine polyacrylamide gel (Bio-Rad) with 100mM Tris 100mM Tricine 0.1% SDS running buffer. GroEL was blotted as a loading control, however uneven transfer at high molecular weights (~60 kDa GroEL vs ~7 kDa CsrA) caused loss of the band in the *Δcsr* lane (confirmed by Coomassie staining of a similar gel after the same transfer protocol). The indicated off-target band suggests consistent loading and was used for normalizing CsrA band quantifications.

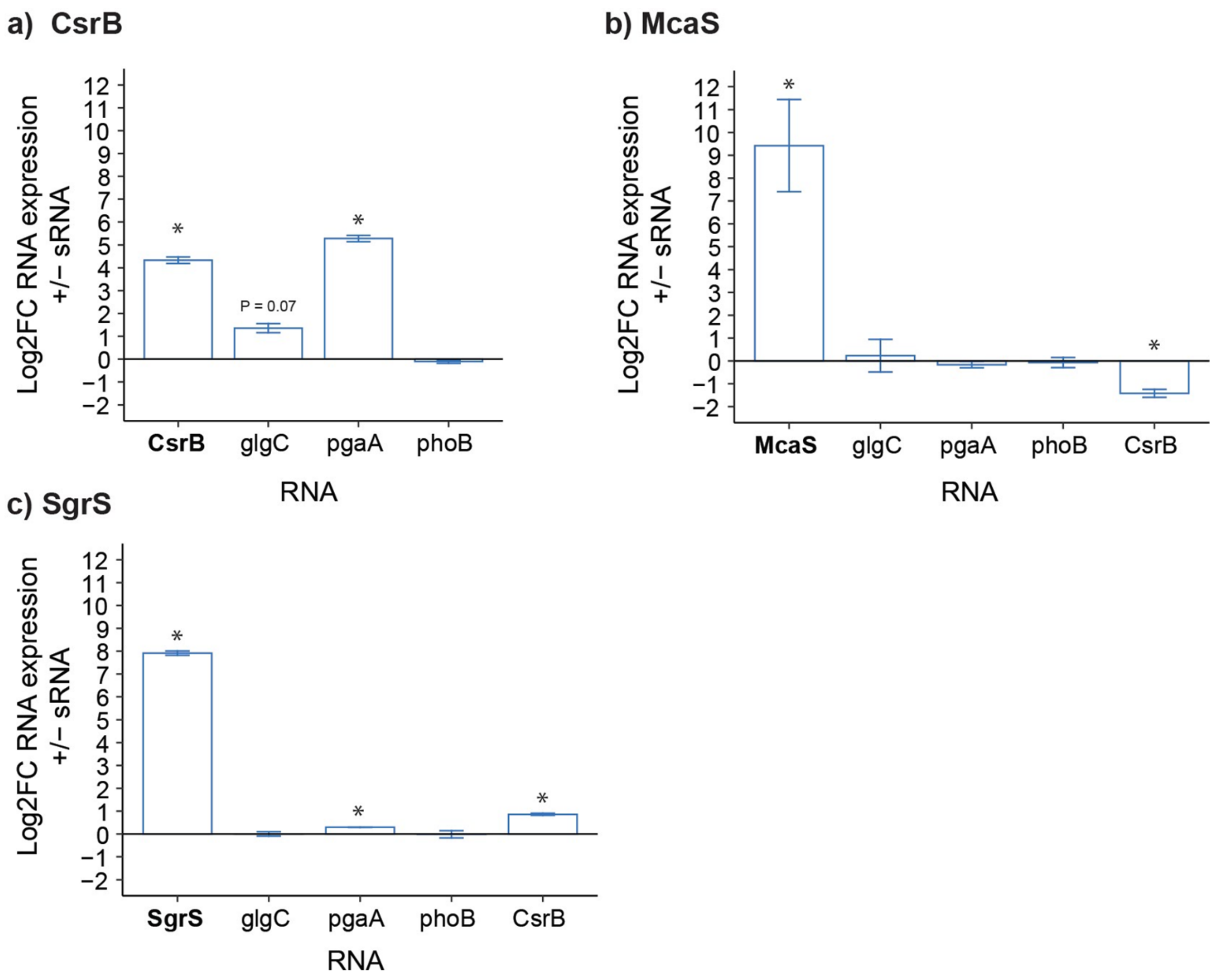


**Supplementary Figure S6. SgrS and McaS do not impact endogenous CsrA-mRNA regulation in mid-exponential phase.** 6a) Log2 fold change (L2FC) of *glgC*, *pgaA*, and *phoB* mRNA abundance after 30 min of CsrB sRNA induction in *ΔcsrBΔcsrC* K-12 MG1655 *E. coli*. CsrB was induced from the from the low copy pBTRK-pLacO plasmid at an $OD_{600}$ of 0.2 and RNA was extracted and quantified by qRT-PCR ($\Delta\Delta C_T$ method) at an $OD_{600}$ of 0.6. Average Log2FC (i.e., - $\Delta\Delta C_T$ values) of 2 or 3 biological replicates are shown with standard deviation error bars. mRNA $C_T$ values are normalized by abundance of the *secA* mRNA (determined with 50 ng RNA per reaction) and CsrB $C_T$ values are normalized by abundance of the 16s rRNA (determined with 0.005 ng RNA per reaction) as described in Material and Methods. Significant results of paired two-tailed t-tests between $\Delta C_T$ values of an RNA in the CsrB induced and CsrB uninduced conditions are reported (P-value < 0.05, asterisks). (6b) Same as in (a), but with McaS sRNA expression in *Δmcas* K-12 MG1655 *E. coli*. (6c) Same as in (a), but with SgrS sRNA expression in *ΔsgrS* K-12 MG1655 *E. coli.*

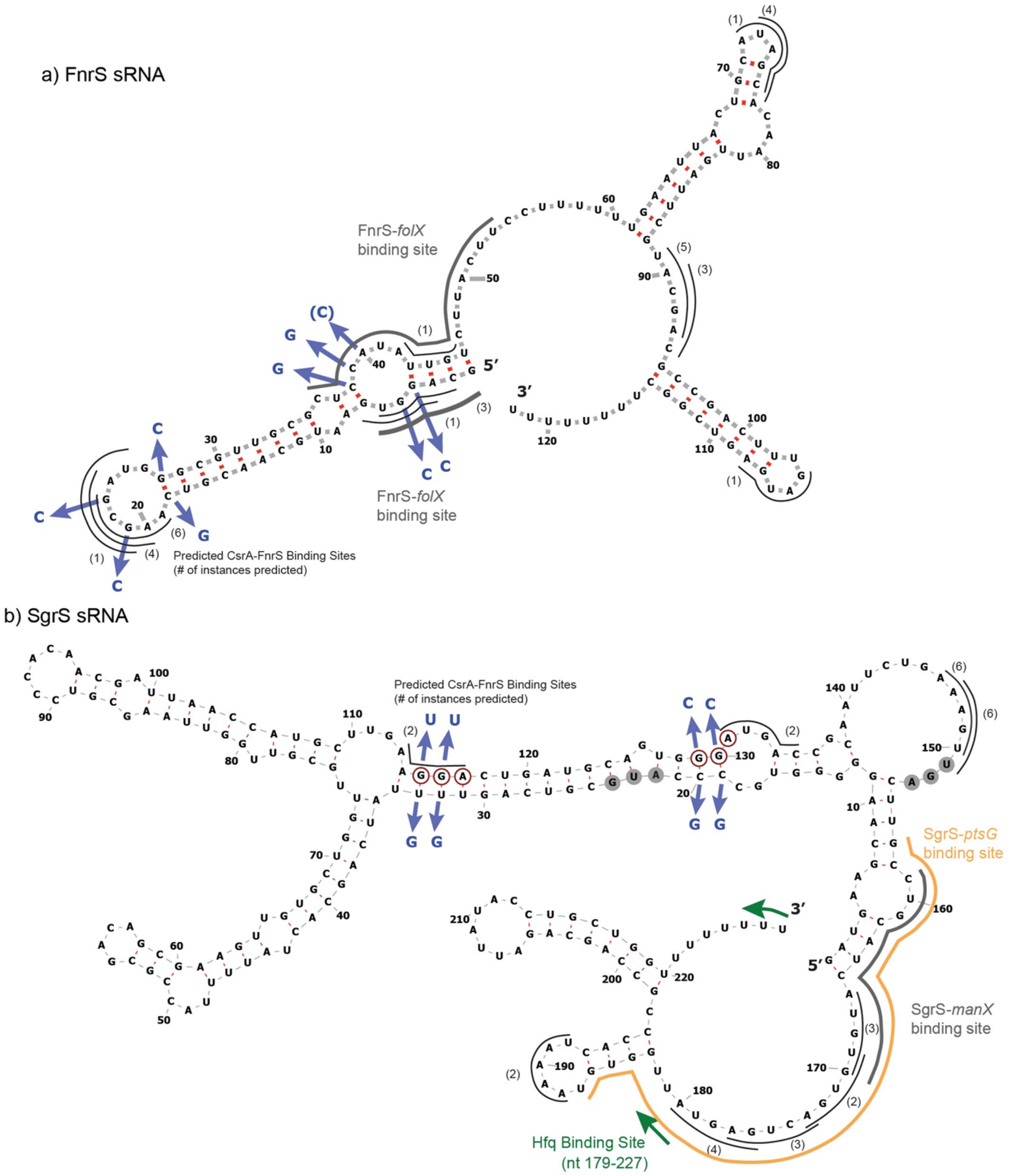


**Supplementary Figure S7. (a) FnrS and (b) SgrS sRNA predicted secondary structures** (Vienna RNA), with GGA motifs (red-circled nts), mutations (blue arrows and nucleotides), and Hfq binding sites (green outlines if known) shown. Predicted CsrA-sRNA binding sites are shown as thin black outlines with the number of times shown in parenthesis that the 5-nt site is predicted among the most-likely 15 pairs of CsrA binding sites. Known mRNA binding sites are indicated as thicker outlines: 7a) FnrS-folX as a gray outline and 7b) SgrS-*manX* and SgrS-*ptsG* as gray and yellow outlines, respectively.

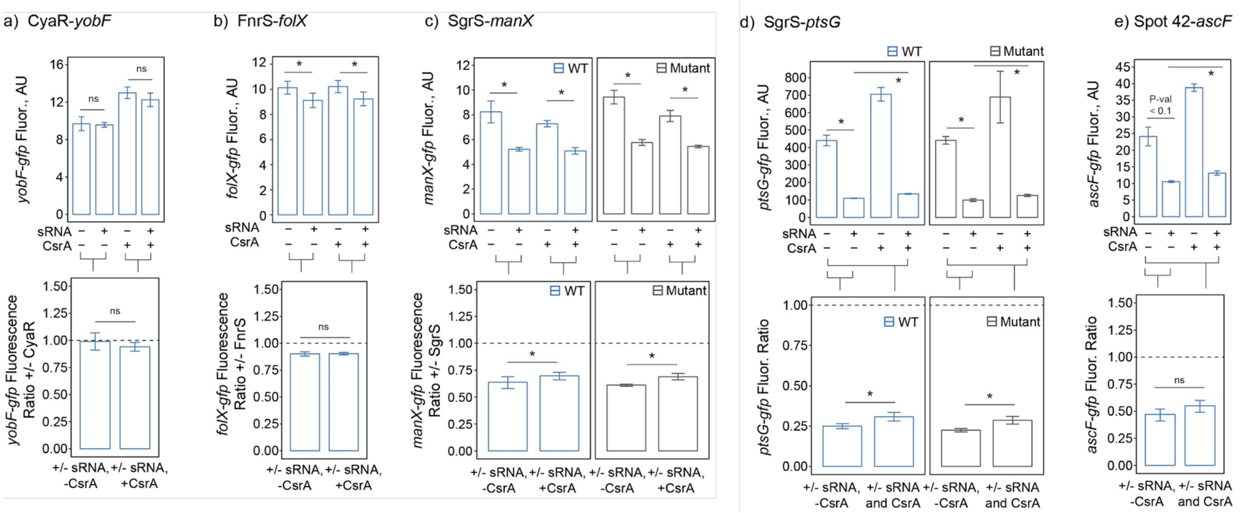


**Supplementary Figure S8. Other sRNA-mRNA regulation screened for CsrA impact.** S8a) CyaR-*yobF* regulation not detected *in vivo*. Fluorescence of the *yobF-gfp* translational reporter at an OD600 of ~0.6 with and without CyaR and CsrA induction (at an OD600 of ~ 0.3) in a pair-wise fashion: –sRNA –CsrA, +sRNA –CsrA, -sRNA + CsrA, and +sRNA +CsrA. Results for wild type CyaR (blue bars) are shown as average median fluorescence values (AU) with standard deviation error (upper panel). Fluorescence ratios of indicated conditions (lower panel). Asterisks indicate P-value < 0.05 of paired, two-tailed t-tests between indicated bars, "ns" indicates not significant. S8b) FnrS-*folX* regulation not impacted by CsrA. Fluorescence of *folX*-*gfp* regulator at an OD600 of ~0.7 with and without FnrS and CsrA induction as in (S8a). Results are shown as in (a). S8c) SgrS-*manX* regulation not impacted by CsrA. Fluorescence of the *manX-gfp* reporter at an OD600 of ~ 0.5 with and without SgrS and CsrA induction as in S8A. Both wild type (blue bars) and GGA mutant (gray bars, see Supplementary Table S1 for sequence) SgrS sRNAs were tested. Results are otherwise as shown as in (a). S8d) SgrS-*ptsG* regulation not clearly impacted by CsrA. Fluorescence of the *ptsG-gfp* reporter at an OD600 of ~ 0.5 with and without SgrS and CsrA induction as in (a). Both wild type (blue bars) and GGA mutant (gray bars, see Supplementary Table S1 for sequence) SgrS sRNAs were tested. Results are otherwise as shown in (a). S8e) Spot 42-*ascF* regulation not clearly impacted by CsrA. Fluorescence of the *ascF-gfp* reporter at an OD600 of ~ 0.7 with and without Spot 42 and CsrA induction as in (a). Results are as shown in (a).

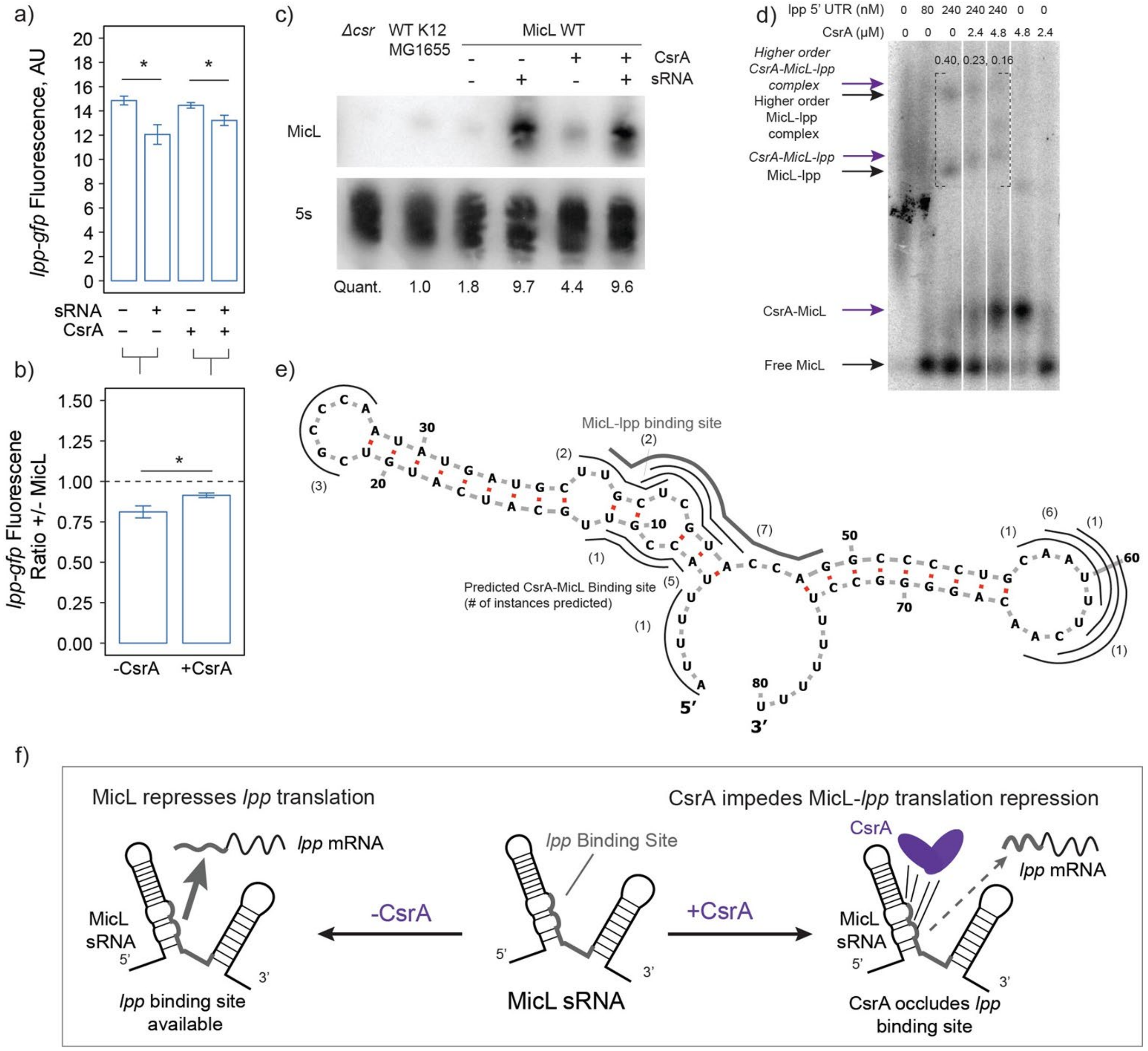


**Supplementary Figure S9. CsrA potentially impedes repression of MicL-*lpp* regulation.** S9a) *In vivo* reporter assay measuring *lpp-gfp* fluorescence in the presence and absence of both the MicL sRNA and CsrA. S9b) Ratios between fluorescence of the *lpp-gfp* reporter with and without MicL expressed evaluated in the presence and absence of CsrA. S9c) Northern blot confirming MicL expression for each induction condition. S9d) EMSA of the MicL sRNA with titrating both CsrA and the *lpp* 5' UTR. When CsrA and *lpp* are present (wells 3-5), ternary complexes appear to form (purple arrows). S9e) Predicted secondary structure of MicL with potential CsrA binding sites predicted (thin black lines) using a biophysical model developed by (Leistra et al., 2017). Confirmed MicL-*lpp* binding sites are noted using thick grey lines. S9f) Potential mechanism of how CsrA remodels MicL-*lpp* interactions by impending translational repression.

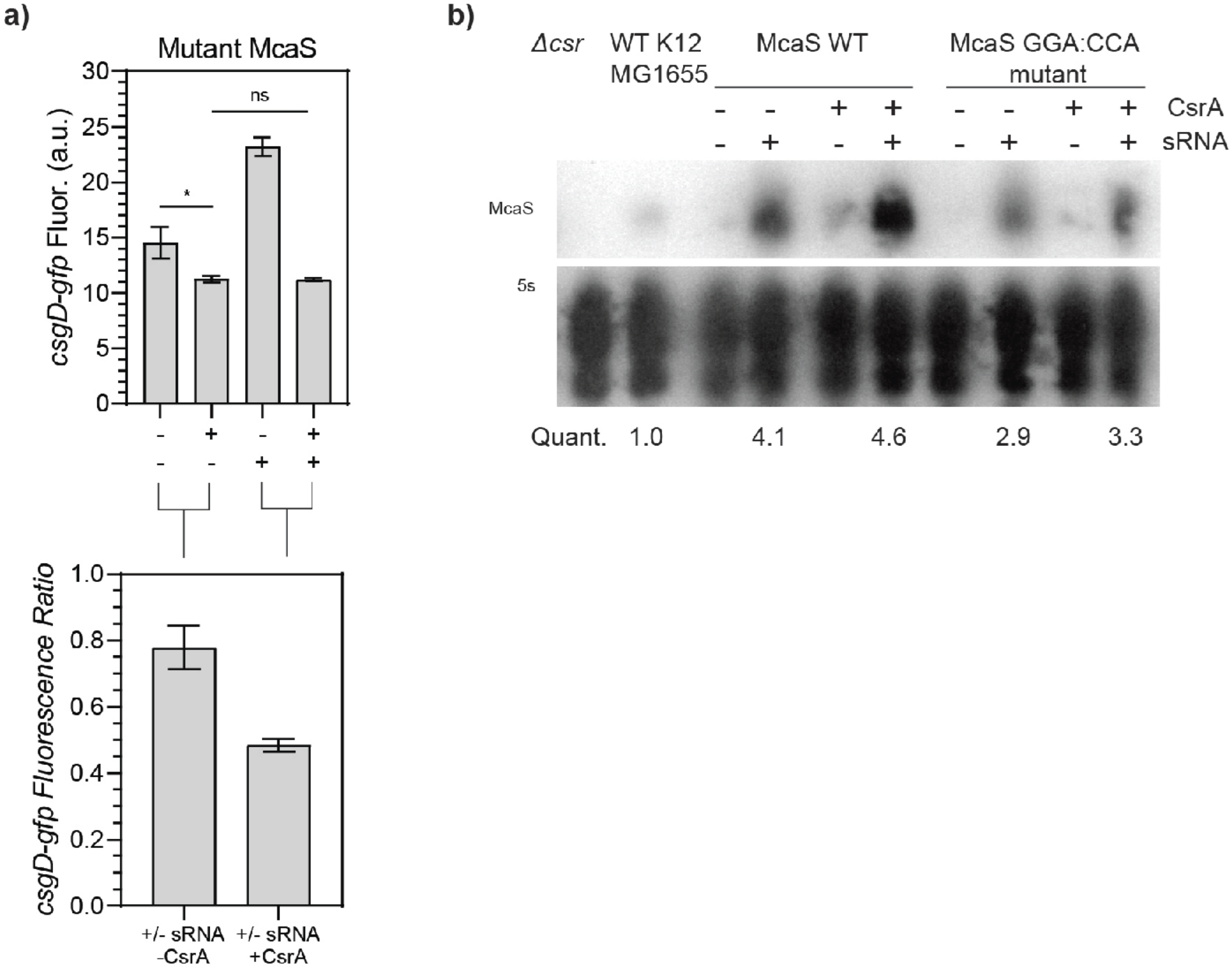


**Supplementary Figure S10. Abrogating CsrA-McaS interactions reduces enhanced repression of McaS-*csgD* by CsrA.** S10a) *In vivo* reporter assay using a mutant McaS, in which the GGA sequence is mutated (Methods and Materials). Eliminated in the CsrA binding site alleviates some of the enhanced repression observed when using the wild type McaS. S10b) Northern blot of the wild type and mutant McaS to show consistent transcript expression under all reporter assay conditions.

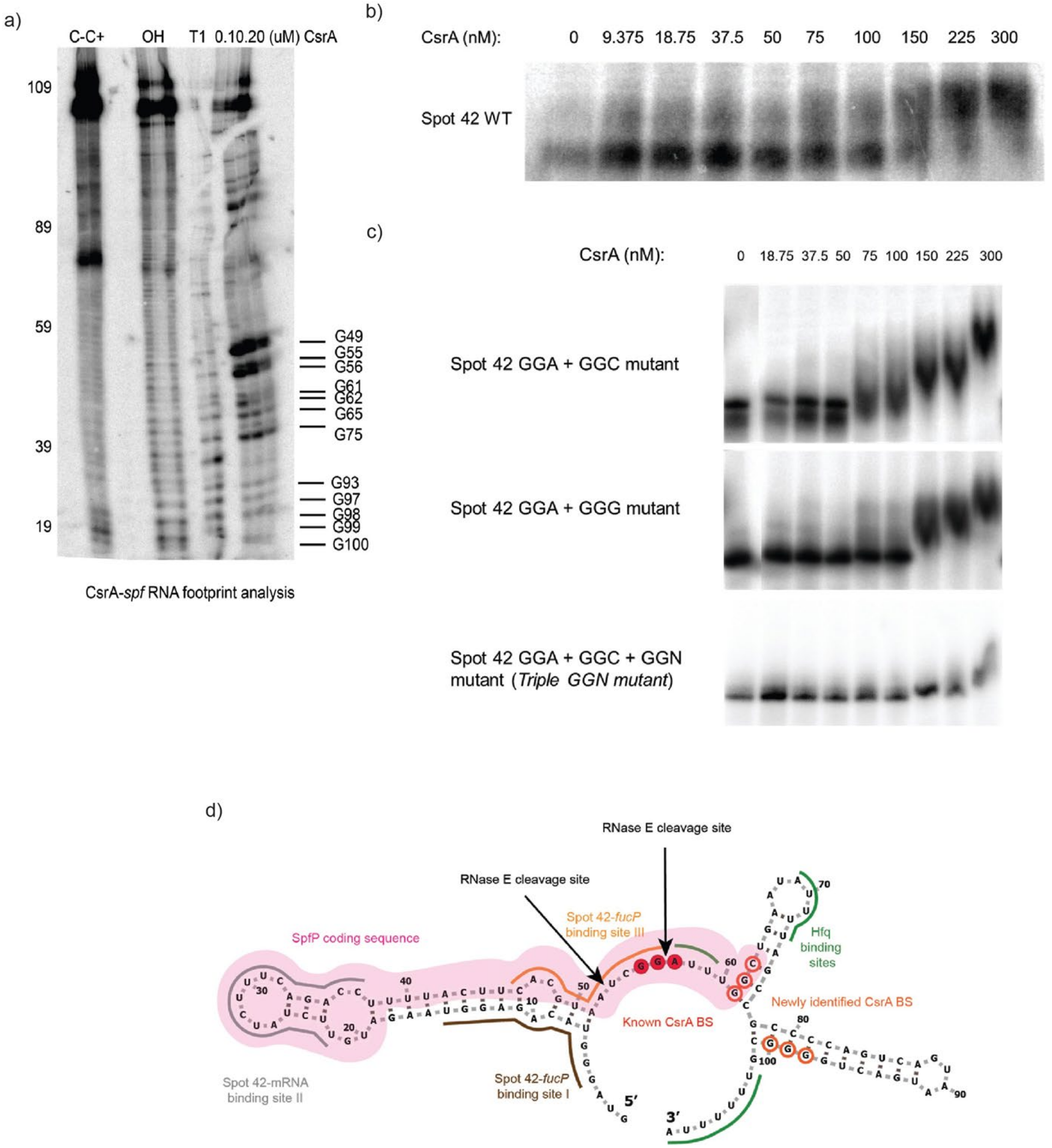


**Supplementary Figure S11. Sequencing gel and EMSAs of Spot42 to identify two novel GGN sequences involved in CsrA-Spot42 binding.** S11a) Sequencing probing gel to evaluate protection of Spot42 under increasing CsrA concentrations (last three lanes). S11b) EMSA of wild type Spot42 sRNA with increasing CsrA concentrations. S11c) EMSAs of Spot42 mutant combinations with increasing concentrations of CsrA. S11d) Predicted secondary structure of Spot42 with known CsrA binding site (dark red circles), novel CsrA binding degenerate GGNs (empty red circles), mRNA binding sites 1-3, the SpfP peptide coding sequence (pink), and Hfq binding region (green).

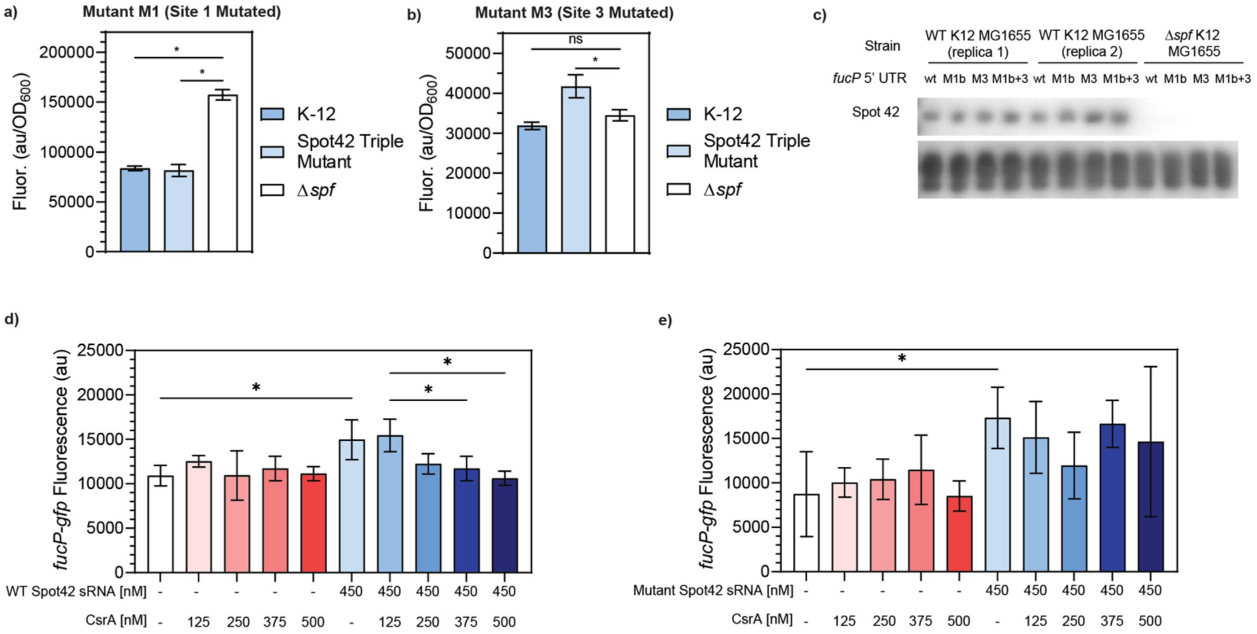


**Supplementary Figure S12. Genomic *in vivo* upon canonical binding site mutation and IVTT evidence that Spot42 can activate *fucP-gfp* expression.** S12a) Expression of *fucP-gfp* transcript with binding site 1 mutated on the *fucP* 5' UTR in *E. coli* expressing wild type Spot42 (K-12) from the genome, a triple mutant Spot42 from the genome (Spot42 Triple Mutant), and *E. coli* with *spf* deleted from the genome. S12b) Expression of the *fucP-gfp* transcript with binding site 3, the canonical Spot42-*fucP* binding site mutated in the same strains of *E. coli*. S12c) Northern blot showing the stability of the Spot42 transcripts expressed from the genome. S12d) IVTT assays measuring the fluorescence of the *fucP-gfp* reporter upon titrating both wild type Spot42 and CsrA. S12e) IVTT assays measuring the fluorescence of the *fucP-gfp* reporter upon titrating both Triple Mutant Spot42 and CsrA.

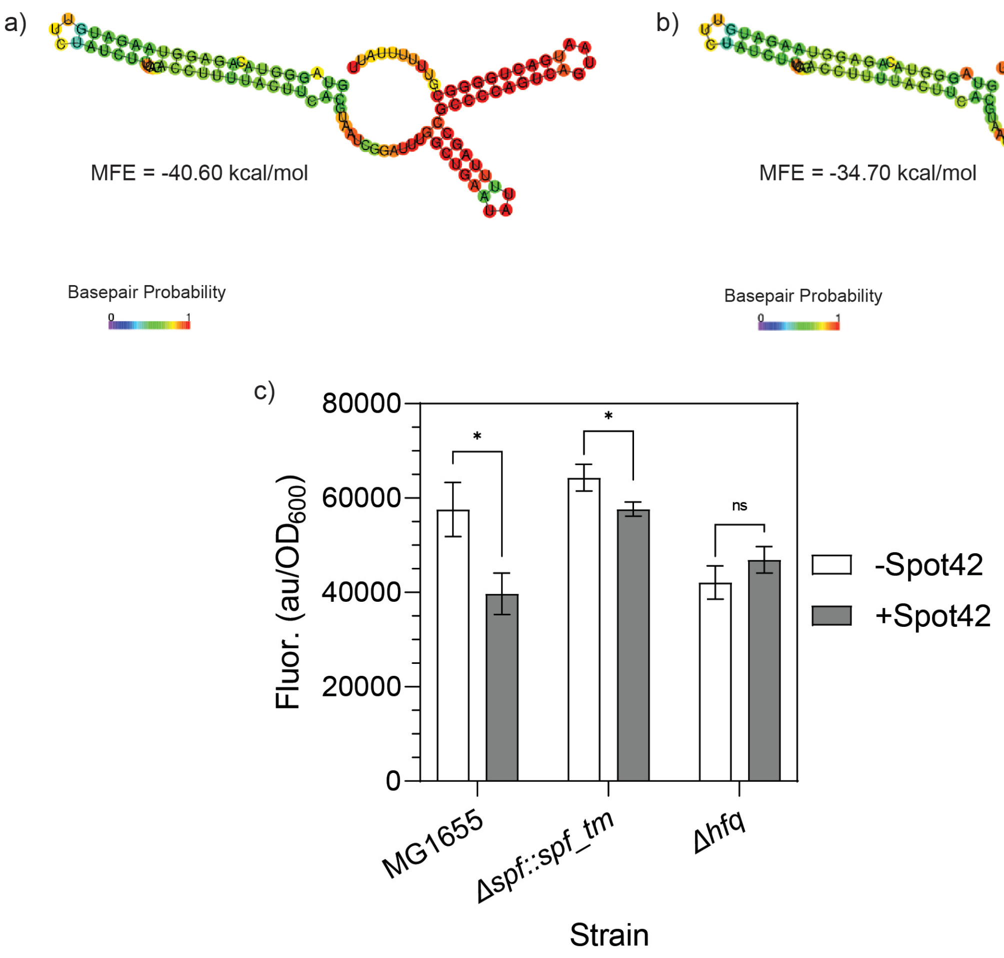


**Supplementary Figure S13. Validation of Spot42 triple mutant secondary structure and mutations do not abrogate native *Hfq-Spot42-fucP* interactions.** S13a) Predicted secondary structure of wild type Spot42 sRNA by the rnafold ViennaRNA Webserver and predicted minimum free energy (MFE). S13b) Predicted secondary structure of the Triple Mutant Spot42 by the rnafold ViennaRNA Webserver and predicted minimum free energy (MFE). S13c) Fluorescence of *fucP-gfp* reporter expressed from the lowcopy pBTRCK plasmid in the presence or absence of Spot42 expressed from the genome following the methods used in Beisel et al. 2011 in wild type *E. coli* (MG1655), a strain with the Spot42 Triple Mutant sRNA integrated to replace the native Spot42 sRNA (*Δspf::spf_tm*), and a *hfq* deletion strain (*Δhfq).* Repression of the *fucP-gfp* reporter is only significantly abrogated in the *Δhfq* strain.

**Summary of Supplementary Tables and Data**

**Supplementary Table S1.** Confirmed mRNA targets and binding sites of sRNAs studied.

**Supplementary Table S2.** Stress association and transcriptional control of sRNAs studied.

**Supplementary Table S3.** Summary of predicted CsrA-sRNA binding sites.

**Supplementary Table S4.** Strains and plasmids used in this study.

**Supplementary Table S5.** Summarized differential expression of mRNA targets of sRNAs studied.

**Supplementary Table S6.** DNA oligonucleotides and fragments used in this study.

**Supplementary Table S7.** Reanalysis of transcriptomics data published in Sowa 2017.

**Supplementary Data S1.** Raw results of Biophysical model-predicted CsrA-sRNA binding sites
